# Genetic Risk and Resilience for Schizophrenia Stratified by Perinatal Gene Expression Predict Adult Cognitive Performance

**DOI:** 10.64898/2026.05.19.726222

**Authors:** Gianluca C. Kikidis, Alessandra Raio, Leonardo Sportelli, Linda A. Antonucci, Alessandro Bertolino, Antonio Rampino, Pierluigi Selvaggi, Daniel R. Weinberger, Giulio Pergola

**Author notes:** **Corresponding author:** Giulio Pergola, Address: Piazza Giulio Cesare, 11 - 70124 Bari, Italy, –.

## Abstract

Genetic risk for schizophrenia (SCZ) has been linked to cognitive performance before the onset age. We examined how SCZ-related polygenic risk and resilience variants, and their co-expression patterns in the human brain, were associated with cognitive abilities across development in 16,520 non-psychiatric European and African ancestry children and adults. SCZ risk showed significant negative associations with spatial, verbal, and working memory across ancestries (all t<-2, pFDR<0.05). In Europeans, risk and resilience variants had opposing effects on attention, working and spatial memory (Δt>4, pFDR<0.05). Polygenic scores filtered through perinatal co-expression networks showed stronger links with cognition than adult (ΔAIC>5.75, p=0.02) or juvenile (ΔAIC>5.8, p=0.03) networks. Cross-ancestry correlations (R=0.52, p<0.01) highlight replicability. These findings support the neurodevelopmental basis of SCZ, suggesting that risk and resilience variants influence cognition from early life, independent of symptoms and elucidate biological pathways through which SCZ risk may influence early cognitive development.

---

Schizophrenia (SCZ) poses a major challenge in psychiatric care, with a lifetime prevalence of 0.5-1.0% and an onset typically occurring between the late teens and mid 20s [1]. Patients with SCZ commonly experience difficulties across multiple cognitive domains, including working memory, executive function, processing speed, attention, and both visual and verbal learning [2–7]. These impairments are well documented to anticipate symptoms and formal diagnosis [8]. Notably, these deficits persist despite currently available pharmacological treatments and have been proposed as indicators of patient outcomes [9].

SCZ is widely recognized to have a genetic component with complex heritability [10]. The latest SCZ genome-wide association study (GWAS) conducted by the Psychiatric Genomics Consortium [11] identified 342 genome-wide significant single-nucleotide polymorphisms (SNPs), accounting for up to 24% of SNP-based heritability (an increase from the 22.5% reported in the previous GWAS [12]). Interestingly, when considering the aggregated effect of SCZ significant genetic variants in a metric also known as polygenic risk score, the majority of individuals within the top percentile for polygenic risk remain unaffected [11]. Hess et al. investigated genetic differences between neurotypical and SCZ-affected individuals with high polygenic risk, identifying variants associated with diminished SCZ liability despite high polygenic risk, termed SCZ resilience [11]. These common genetic variants are thought to promote resilience by decreasing the penetrance of SCZ risk loci. In fact, their findings suggest that in affected individuals, genetic risk and resilience variants, located at distinct loci, are correlated, in contrast to healthy individuals. This may suggest that a greater genetic risk is required to overcome resilience in individuals who develop SCZ. Moreover, recent studies have shown that polygenic resilience scores positively correlate with cognitive performance in individuals at risk for psychosis [12], in contrast with polygenic risk scores which tend to show the opposite correlation with cognition. This evidence points to a potentially protective effect of cognition on SCZ liability and suggests core pathways and processes in which SCZ risk and resilience converge.

A growing body of evidence links SCZ risk variants to alterations in cognitive phenotypes [13, 14], emphasizing the relationship between SCZ and neurocognitive systems [9, 15, 16]. Genetic factors contributing to cognitive abilities are shared with those linked to SCZ [17, 18] and are heritable, as shown by twin studies [19] and studies involving relatives of SCZ patients [15, 20]. Cognition is reported to account for at least one-third of the variance explained by genetic risk factors for SCZ [21]. Indeed, polygenic scores (PGSs) for SCZ risk are negatively associated with cognitive performance, even in healthy controls [22]. These findings tie in with the idea that SCZ mechanisms in brain manifest before clinical diagnosis, affecting brain development associated with poor cognitive function [23]. It has been proposed that the biological substrate that underlies the emergence of the clinical syndrome of schizophrenia in early adulthood shows alternative phenotypic expression based on a developmental trajectory that changes over time [24]. While psychosis per se is a rare characteristic of brain dysfunction before early adolescence, cognitive dysfunction can manifest much earlier. Cognitive performance may thus reflect an early manifestation of illness biology, a prodrome, a harbinger in its own right of genetic and environmental risk. Previous reports have shown that relatively increased genetic risk for SCZ is also associated with poor cognitive function emerging well before the typical age of onset of SCZ (e.g., before 10 years old), particularly in domains such as emotion recognition speed [26], verbal reasoning [26], and working memory [2]. Disentangling the role of cognitive development in SCZ requires a biological understanding of the shared genetic factors behind their relationship.

Neurodevelopmental trajectories related to SCZ risk manifest as early as the perinatal stage of life [23], e.g., as cognitive and social deficits that persist from childhood to adulthood [3], when they influence vulnerable neuronal systems—such as dopaminergic pathways affecting the prefrontal cortex—that mature during adolescence or early adulthood. Consistently, the neurodevelopmental hypothesis of SCZ posits that pathogenic processes occurring early in life that underlie the eventual expression of psychosis, particularly those affecting the relatively late development and function of the dorsolateral prefrontal cortex, remain clinically silent until they interact with neurodevelopmental processes that occur much later. These interactions coincide with the maturation of key processes underlying the main symptoms of the disorder [24, 27–29]. Recent advances in transcriptomics across development have supported this view [30–34]. The biological basis of vulnerability may, for instance, take the form of altered circuits of neurotransmission activity that are susceptible to stress and may promote the clinical manifestation of psychotic symptoms [35].

The dynamic nature of gene expression across development is an emerging factor in understanding the etiology of complex developmental brain disorders [36]. Polygenic scores (PGSs) estimate an individual’s genetic predisposition to various conditions without accounting for the time when genetic predispositions are expressed, thus overlooking the age-specific relevance of the implicated biological processes [37–39]. As shown by Van Der Meer et al. [40], PGSs parsed by temporal expression patterns explain variance in SCZ diagnosis differently based on prenatal and postnatal gene expression, while also interacting with each other. Gene expression profiles and their co-expression contexts shift significantly across developmental stages [41, 42]: certain genes become more active or change their expression partners at specific stages due to developmental roles. Weighted Gene Co-expression Network Analysis (WGCNA) [30] has been used to identify clusters of co-expressed genes, called modules, in post-mortem brain tissues specific to different life stages, e.g., perinatal, juvenile, and adult periods. Studying gene co-expression patterns and genetic variation in an age-specific context can disentangle the polygenic risk of living individuals into these distinct developmental stages, allowing insights into how co-expression networks evolve over time. This approach, in turn, allows the study of molecular contexts of risk unfolding along neurodevelopment.

Although PGSs do not reflect biology per se, as they simply consist of allele frequency differences between cases and controls, they can be parsed into gene sets that reflect risk within certain molecular contexts. Parsing genetic risk based on co-expression patterns across developmental stages thus confers a functional and temporal biological dimension to PGSs [43]. Intriguingly, since all adults were once infants, PGSs parsed for perinatal biology remain informative even when applied to adult samples. Using perinatal-based scores in adults has the potential to help identify individuals at higher risk due to early developmental factors and distinguish them from individuals carrying more risk in biological pathways expressed in later life stages. Such heterogeneity of risk may be linked to clinical manifestations, e.g., striatal dopamine synthesis [45].

Importantly, most existing genetic studies of SCZ and cognitive function have been conducted in individuals of European ancestry, limiting the generalizability of findings across diverse populations [46, 47]. Individuals of African ancestry are underrepresented in psychiatric genomics despite their greater genetic diversity and specific linkage disequilibrium patterns, which provide both challenges and unique opportunities for discovering novel biology [48–50]. Here, we investigate the potential influence of polygenic risk scores on neurocognitive performance across different age periods in neurotypical individuals of European and African ancestries and the diverging effects of polygenic risk and resilience scores in a European subgroup. Genetic resilience scores reflect common variants that mitigate SCZ risk. However, genetic resilience summary statistics are not currently available for non-European populations. Thus, resilience analyses were restricted to individuals of European descent. We hypothesized that, if risk and resilience represent genetically different entry points into neurobiological processes core to SCZ, then they should impact similar cognitive domains in opposite directions. To account for known sex-related differences in cognitive performance, we also explored potential differences in neurocognitive associations with genetic risk between male and female individuals. Moreover, we investigated as a positive control the impact of genetic variants for cognitive and general intelligence on cognitive performance in contrast to the negative effect caused by SCZ-related variants. Finally, we investigated the effects of specific polygenic variants parsed by age period-specific co-expression patterns in post-mortem human brain tissue on cognitive functions to better characterize the impact of biological pathways involved in distinct life stages.

## Results

### SCZ risk and resilience PGS association with cognitive performance in children, adolescents, and adults

We examined the effect size of risk PGS on each cognitive domain within each cohort for both European (EUR) and African (AA) ancestry with a linear regression model to explore the impact of SCZ genetic risk on cognitive performance. We found a significant negative association in both EUR and AA individuals in spatial memory (NIH_EUR_ t = -3.4, p = 0.008, NIH_AA_ t = -2.25, p = 0.02), working memory (ABCD_EUR_ t = -3, p = 0.009, ABCD_AA_ t = -2.35, p = 0.01), verbal memory (PNC_EUR_ t = -3.01, p = 0.008, PNC_AA_ t = -2.08, p = 0.03), and cognitive flexibility (ABCD_EUR_ t = -2.2, p = 0.03, ABCD_AA_ t = -2.45, p = 0.01). We also found significant negative associations in EUR individuals in working memory (PNC t = -2.5, p = 0.02) and cognitive flexibility (PNC t = -2.42, p = 0.02), and in AA individuals in spatial memory (NIH t = -2.23, p = 0.02) and spatial reasoning (PNC t = -2.1, p = 0.03). All results are shown in Figure 1a.

**Figure 1:**
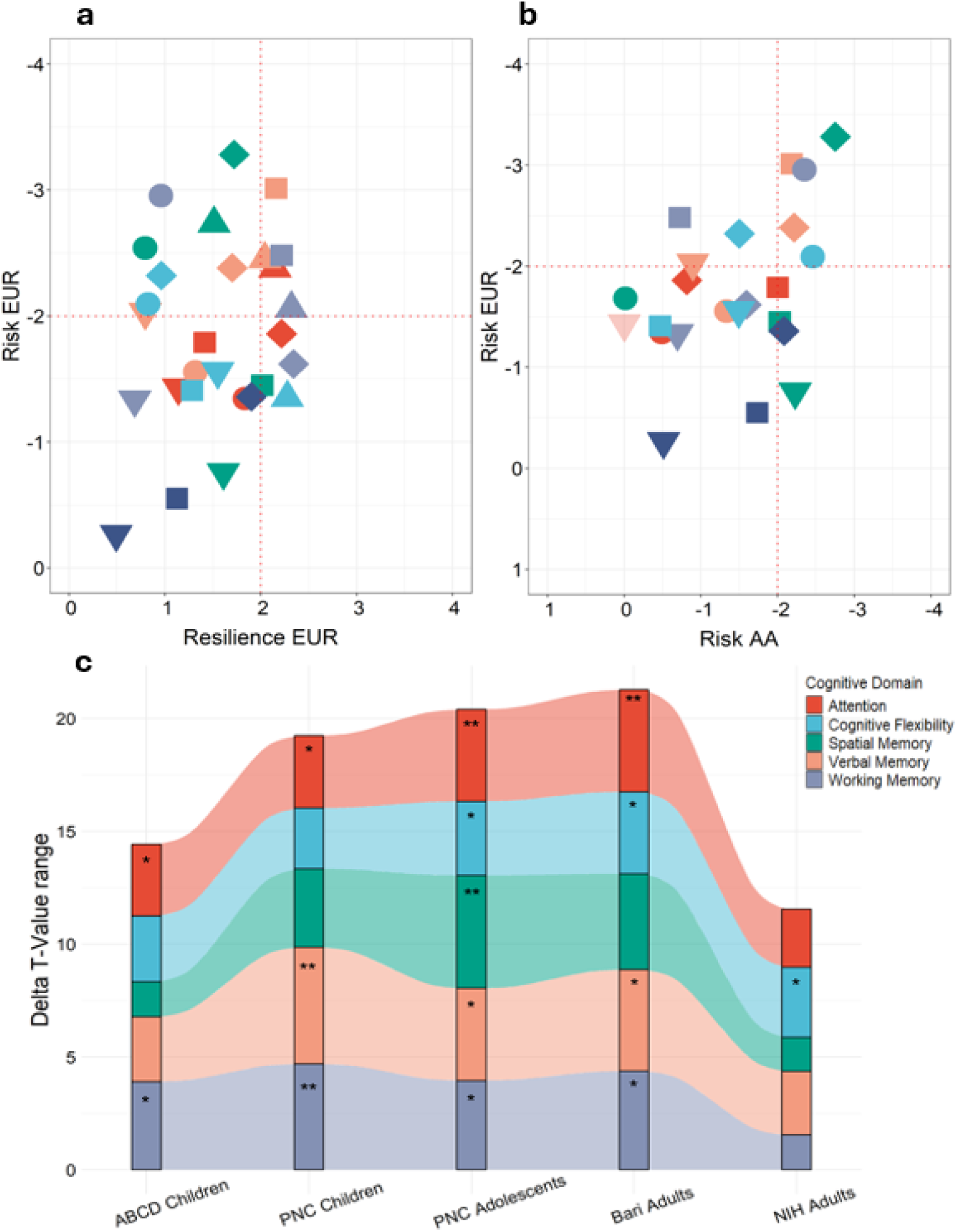
**a**) Risk t-values for each cognitive domain and dataset for both European (EUR) and African (AA) ancestry. Red lines indicate significance as follows: upper left quadrant p < 0.05 for EUR samples, lower right quadrant p < 0.05 for AA samples, upper right quadrant p < 0.05 for both EUR and AA samples. **b**) Risk and resilience t-values for each cognitive domain and each dataset. Red lines indicate significance as follows: lower right quadrant p<0.05 for resilience scores, upper left quadrant p<0.05 for risk scores and upper right quadrant p<0.05 for both risk and resilience scores. **c)** Risk and resilience Delta PGS effects in the ABCD, PNC, Bari and NIH cohorts on all cognitive domains. Delta for T values for each score are shown on the y-axis. Associations are statistically significant in attention, cognitive flexibility, spatial memory, verbal memory and working memory performance. Asterisks signal the significance of linear hypothesis tests between risk and resilience estimates (ns: p>0.05, *: p<=0.05, **: p<=0.01, ***: p<=0.001). Connections between bar plots highlight the changes in effect sizes between different cohorts ordered by median age from left to right.

We further examined the disparity in effect sizes between risk and resilience on each cognitive performance test within each cohort by calculating the t-value difference between each score, aiming to elucidate the potential divergent effects of the genetic components on cognitive performance (Figure 1 b and c). This approach is equivalent to comparing the distribution of scores against each other rather than against zero. Delta values represent the absolute difference between risk and resilience effect sizes in terms of t-values. We found significant divergence in children and adults in cognitive flexibility (PNC Δ t-value: 4.8, linear hypothesis test (LHT) p_FDR_ = 0.02; Bari Δ t-value: 3.71, LHT p_FDR_ = 0.04), attention (ABCD Δ t-value: 3.17, LHT p_FDR_ = 0.04; PNC Δ t-value: 3.68, LHT p_FDR_ = 0.03; Bari Δ t-value: 4.09, LHT p_FDR_ = 0.007; NIH Δ t-value: 2.57, LHT p_FDR_ = 0.20), spatial memory (PNC Δ t-value: 4.89, LHT p_FDR_ = 0.002; Bari Δ t-value: 4.24, LHT p_FDR_ = 0.003), verbal memory (PNC Δ t-value: 3.89, LHT p_FDR_ = 0.02; Bari Δ t-value: 4.36, LHT p_FDR_ = 0.01) and working memory (ABCD Δ t-value: 3.91, LHT p_FDR_ = 0.03; PNC Δ t-value: 4.72, LHT p_FDR_ = 0.01; Bari Δ t-value: 4.23, LHT p_FDR_ = 0.01). These differences, represented by Delta T-values, are visually depicted in Figure 1b. Risk and resilience individual t-values showed a mean correlation of Pearson’s r_g_ = -0.65 across all cognitive domains, although the scores were not correlated in the samples tested (mean Pearson’s r_g_ = 0.03), as also in Hess et al., 2020 [51]. The metanalysis across all cohorts showed significant FDR-corrected meta p-values for attention (risk: β = -0.05, 95% CI [-0.08, -0.02], p = 8.3×10^−4^, resilience: β = 0.05, 95% CI [0.03, 0.08], p = 9.4×10^−4^), working memory (risk: β = -0.06, 95% CI [-0.09, -0.04], p = 6.6×10^−5^, resilience: β = 0.06, 95% CI [0.02, 0.09], p = 3.2×10^−3^), and verbal memory (risk: β = -0.07, 95% CI [-0.10, -0.04], p = 4.0×10^−6^, resilience: β = 0.05, 95% CI [0.02, 0.07], p = 3.6×10^−3^).

To offer a fair comparison of sample sizes between risk and resilience GWAS, we additionally calculated the risk PGS based on the same PGC wave [12] used for the resilience GWAS and calculated the Δ T-values with the resilience PGS effects. The resulting associations were comparable to our results (Pearson’s r = 0.88), suggesting that sample size discrepancies do not play a major role in our pattern of results (Supplementary Figure 1).

There were no significant interactions between risk and resilience scores in association with any cognitive domain after multiple comparisons correction (Supplementary Figure 2). Therefore, risk and resilience associations indeed converge on cognitive domains despite tapping onto different genetic variants, in absence of significant two-way interactions.

### SCZ risk and resilience PGS interaction with age and genetic sex

Nominally significant interactions between risk, resilience, and age are reported in Supplementary Figure 3. None survived false discovery rate (FDR) correction. Similarly, no interactions between risk and resilience PGSs and biological sex survived FDR correction (see Supplementary Figure 4).

### Cognitive and general intelligence PGSs association with cognitive performance in children, adolescents, and adults

In addition to SCZ risk and resilience, in our children and adolescents’ samples, variants associated with cognitive intelligence, i.e., variants found significant in a general cognitive function component derived from different cognitive tasks including at least three different cognitive domains [17]were strongly associated with all cognitive domains in children and adolescents from the ABCD and PNC cohorts, while revealing a significant effect on adults only in verbal memory. Furthermore, general intelligence, i.e., variants linked to variation in intelligence assessed via different cognitive tasks, mainly testing fluid domains of cognitive functioning [18]was associated positively with working memory, spatial memory and verbal memory in children and adolescents. All effect sizes are shown in Table 2 and results are illustrated in Figure 2. The age pattern of effect sizes is evidently different from the case of SCZ risk and resilience shown in Figure 1.

**Figure 2:**
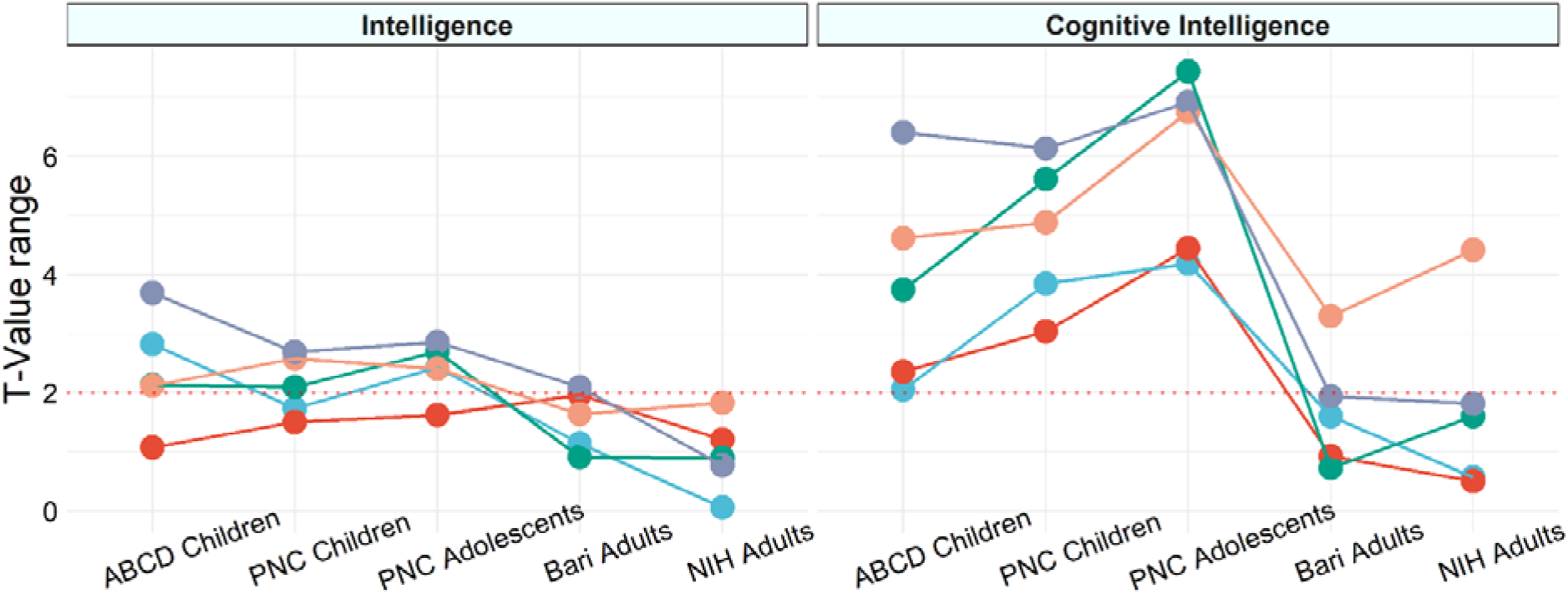
cognitive performance and PGS for cognitive (Figure 2 a) and general (Figure 2 b) intelligence in ABCD, PNC, Bari and NIH. T values for all PGSs are shown on the y-axis. Associations are statistically significant in attention, cognitive flexibility, spatial memory, verbal memory and working memory performances. Connections between dot plots highlight the changes in effect sizes between different cohorts ordered by median age from left to right.

### Other disorders

We identified significant (all p_FDR_ < 0.05) negative associations between risk for ADHD and attention in PNC (t-value: -2.7) and working memory in ABCD (t-value: -4.3). MDD risk was negatively associated with attention and spatial reasoning in the PNC sample (t-value: -3.2, -4.0 respectively) and with working memory, cognitive flexibility and spatial memory in the NIH sample (t-value: -2.5, -3.6, -7.1, respectively). No other psychiatric, neurodegenerative or non-neurological genetic variation showed significant associations with cognitive performance. Results for all disorders are shown in Supplementary Figure 5. ADHD and MDD risk appear to negatively influence cognitive performance across multiple domains, in contrast to general intelligence and cognitive function polygenic scores.

### SCZ risk and resilience PGSs association with cognitive performance parsed for biological pathways and age stages derived from post-mortem samples

To assess model fit, we used Akaike Information Criterion (AIC)—an index that penalizes high predictor dimensionality while estimating the goodness of fit. We first looked at age-parsed WGCNA risk modules in both EUR and AA individuals and found a better model fit in perinatal co-expression modules for attention (Bari_EUR_ Δ*_AIC=_ 1.42*, ABCD_AA_ Δ*_AIC=_* 1.02), cognitive flexibility (PNC_AA_ Δ*_AIC=_ 1.45*, NIH_AA_ Δ*_AIC=_ 1.62*), spatial memory (NIH_EUR_ Δ*_AIC=_ 2.84*), verbal memory (PNC_EUR_ Δ*_AIC=_ 1.09, ABCD_AA_* Δ*_AIC=_ 1.42*), and working memory (PNC_AA_ Δ*_AIC=_ 2.22)*. In the European subsample, age-parsed WGCNA modules grouped by perinatal age stage showed a better model fit compared to all risk and resilience variants PGSs with a significantly lower AIC in adults for spatial memory (Bari Δ*_AIC=_* 2.2), verbal memory (Bari Δ*_AIC=_* 1.8), and attention (Bari Δ*_AIC=_* 1.7, NIH Δ*_AIC=_* 1.7). Additionally, children and adolescents showed improved model accuracy with perinatal parsed modules in spatial reasoning (PNC adolescents Δ*_AIC=_* 1.8, PNC children Δ*_AIC_* = 1.6), working memory (PNC children Δ*_AIC=_* 1.6), and verbal memory (PNC children Δ*_AIC=_* 1.6, PNC adolescents Δ*_AIC=_* 1.3). These results are shown in Figure 3 and suggest that gene co-expression networks parsed by developmental age stages, specifically perinatal co-expression, offer a more granular framework for modeling cognitive performance than traditional PGS approaches in children and adults. All correlations between observed and predicted cognitive performances with standard PGSs and age-parsed PGSs for SCZ risk and resilience variants are reported in Supplementary Figure 6.

**Figure 3:**
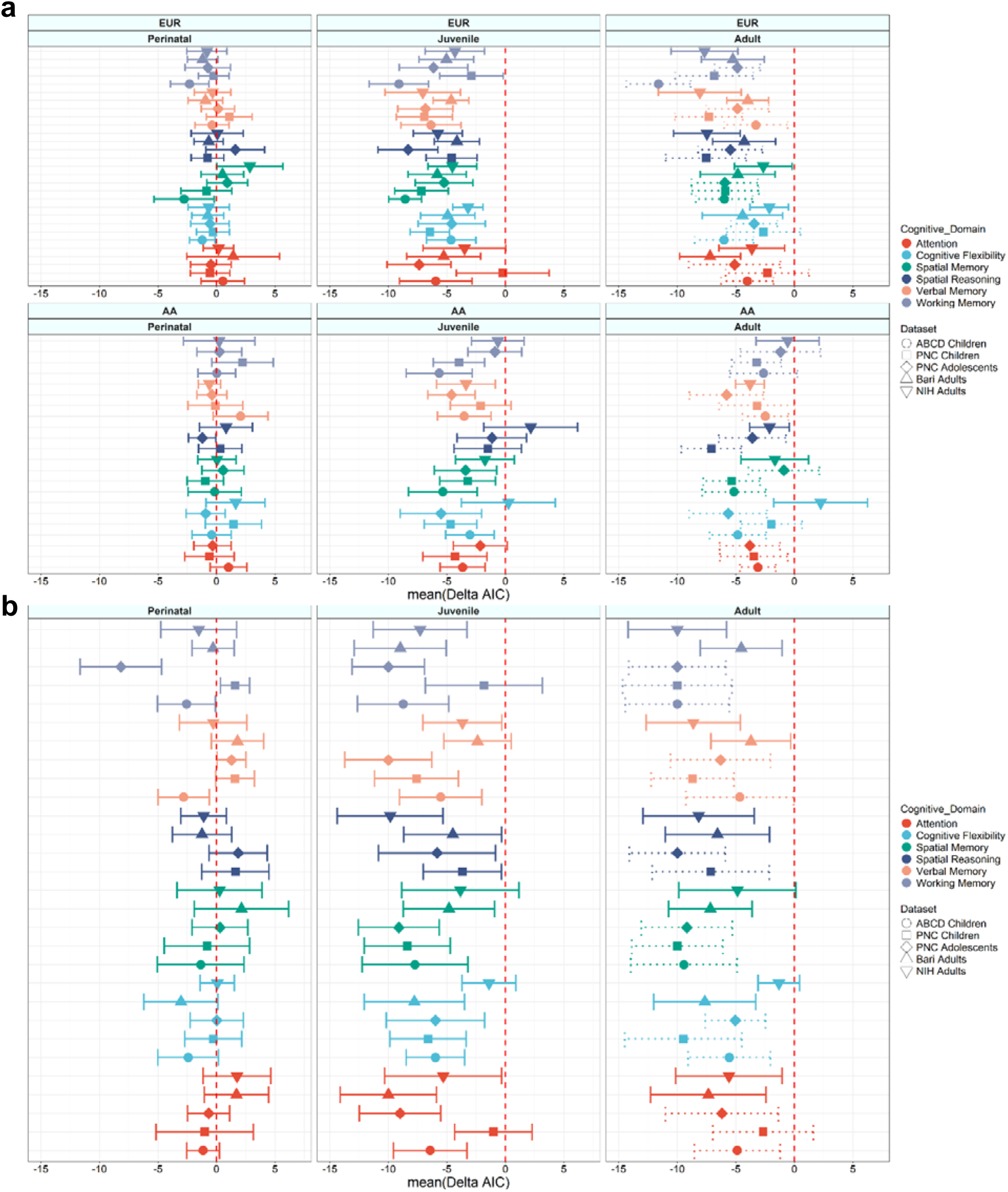
**a)** Median Δ_AIC_ performance between WGCNA risk modules in each cohort and cognitive domain for both EUR and AA samples. Positive Δ_AIC_ values indicate lower AIC for WGCNA modules, meaning that the model performance is higher compared to the whole PGScs model. Error bars indicate standard error across 100 iterations. Asterisks indicate Wilcoxon paired t-test significance (* p < 0.05, ** p < 0.01, *** p < 0.001). Adult parsing in ABCD and PNC is shown in dotted lines, since these networks did not yet take place in children and adolescents and act as a negative control. **b)** Median Δ_AIC_ performance between WGCNA risk and resilience modules grouped by age periods and whole PGScs for all cognitive domains in the EUR cohorts.

**Figure 4:**
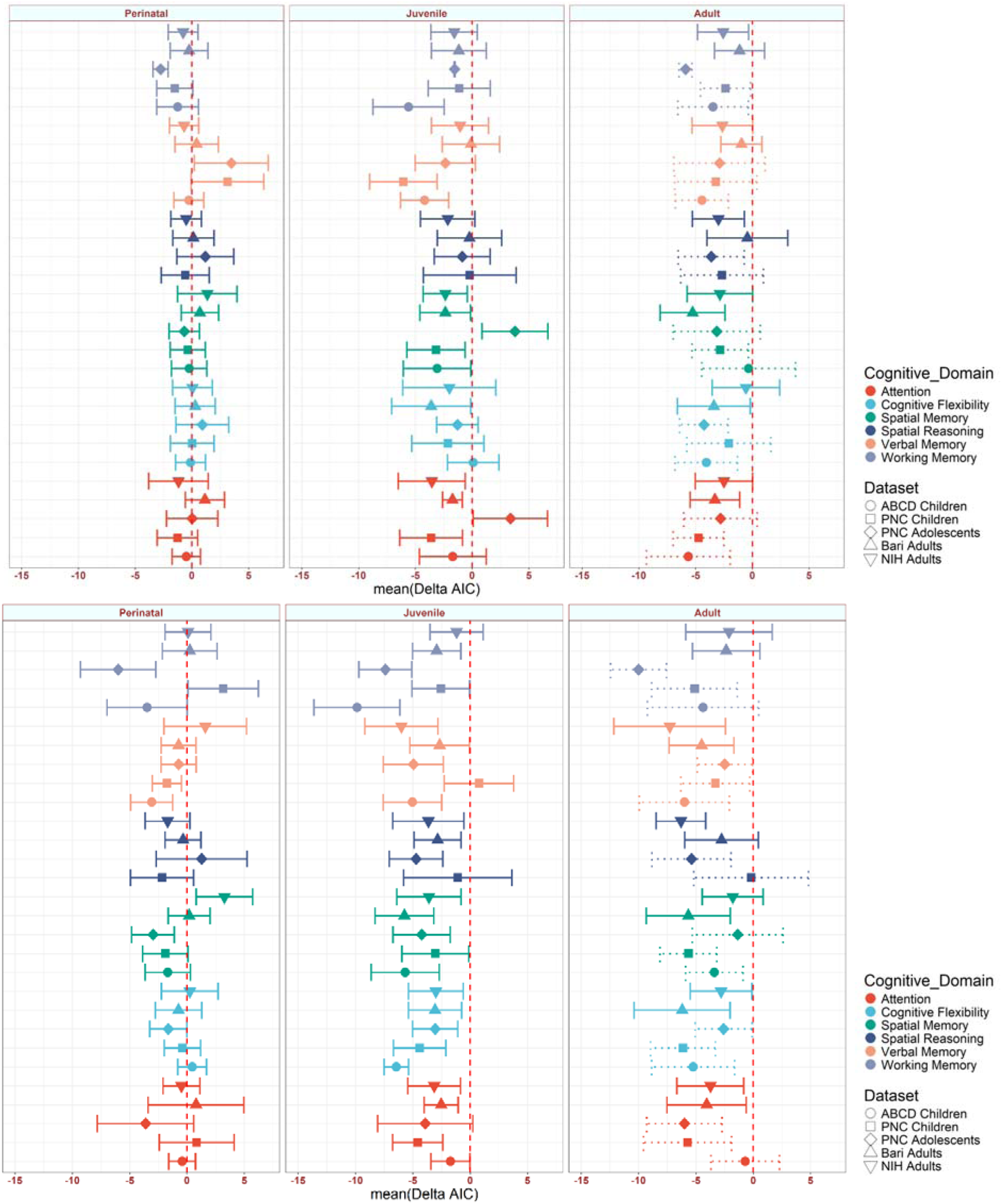
Median Δ_AIC_ performance between WGCNA modules grouped by age periods and whole PGScs for all cognitive domains and all cohorts. Asterisks indicate Wilcoxon paired t-test significance (* p < 0.05, ** p < 0.01, *** p < 0.001). Adult parsing in ABCD and PNC are shown as dotted lines. Figure 4 a represents cognitive intelligence and Figure 4 b represents general intelligence.

Across cognitive domains, gene co-expression networks defined by perinatal risk exhibited better fits, on average, relative to both adult (EUR: Δ_AIC_ = 5.09, p_wilcoxon_ = .02; AA: Δ_AIC_ = 3.37, p_wilcoxon_ = .03) and juvenile (EUR: Δ_AIC_ = 5.25, p_wilcoxon_ = .02; AA: Δ_AIC_ = 3.04, p_wilcoxon_ = .03) risk-derived networks. Notably, adult and juvenile co-expression networks did not differ significantly in their association with cognitive outcomes (AA: Δ_AIC_ = 0.33, p_wilcoxon_ = 1; EUR: Δ_AIC_ = 0.16 p_wilcoxon_ = 1). Cross-ancestry analysis revealed a moderate correlation (Pearson’s R = 0.52, p < 0.01), further underscoring the distinctive relevance of the perinatal molecular landscape in shaping associations with later-life cognitive phenotypes.

In analyses restricted to the European-ancestry subsample, inclusion of resilience-associated co-expression modules did not alter the primary finding: perinatal risk-derived co-expression networks continued to outperform both adult (Δ_AIC_ = 6.63, p_wilcoxon_ = .01) and juvenile (Δ_AIC_ = 5.88, p_wilcoxon_ = .02) counterparts.

### General and cognitive intelligence PGSs association with cognitive performance parsed for biological pathways and age stages derived from post-mortem samples

Age-parsed WGCNA modules for cognitive intelligence grouped by perinatal age stage showed a better model fit compared to all variant PGS in adults for spatial (NIH Δ*_AIC=_* 3.25) and verbal memory (NIH Δ*_AIC=_* 1.60), and in children for working memory (PNC children Δ*_AIC=_* 3.20) and spatial reasoning (PNC adolescents Δ*_AIC=_* 1.27). Perinatal co-expression modules for general intelligence showed a lower AIC in children within the domains of verbal memory (PNC children Δ*_AIC=_* 3.20, PNC adolescents Δ*_AIC=_* 3.47) and spatial reasoning (PNC adolescents Δ*_AIC=_* 1.18), and in adults within spatial memory (NIH Δ*_AIC=_* 1.35) and attention (Bari Δ*_AIC=_* 1.14). Juvenile co-expression modules were significant in PNC adolescents in the domains of spatial memory (Δ*_AIC=_* 3.77) and attention (Δ*_AIC=_* 3.37). Supplementary Figures 7 and 8 illustrate the correlations between observed and predicted cognitive performances with standard PGSs and age-parsed PGSs for cognitive and general intelligence variants in the European subsample.

The perinatal co-expression PGSs explained cognition slightly better compared to adult (Δ*_AIC_*= 3.50, p_wilcoxon_ = .05) and juvenile (Δ*_AIC_* = 3.12, p_wilcoxon_ = 0.06) co-expression. Adult and juvenile co-expression patterns were again similar (Δ*_AIC_* = 0.38, p_wilcoxon_ = 1). General intelligence co-expression differences were similarly low, with perinatal co-expression performing slightly better than adult (Δ*_AIC_* = 3.00, p_wilcoxon_ = .09) co-expression, while juvenile and adult co-expression models were not significantly different (Δ*_AIC_* = 1.22, p_wilcoxon_ = 1).

### Gene ontology enrichment analysis

We performed gene ontology (GO) enrichment analysis to investigate the enrichment of significant predictive genes across age stages for biological pathways. In juveniles (combined ABCD and PNC cohorts), cognitive flexibility was significantly associated with synaptic and developmental pathways. The age-specific PGSs showed a consistent pattern, privileging the juvenile co-expression networks. Verbal memory was also significantly associated with synaptic pathways in juvenile co-expression networks. Findings between European and African children differed, possibly because different co-expression modules were selected within the algorithm based on gene proximity to different variants. Nevertheless, biological functions such as cell adhesion and transport were replicable across ancestries and were associated with the same cognitive domains. In adults (combined Bari and NIH cohorts), working memory and attention were significantly associated with synaptic pathways within the juvenile co-expression, while spatial memory and cognitive flexibility were associated with myelination, metabolism, and membrane-related biological processes in the perinatal co-expression networks. Figure 5 illustrates these results, suggesting that age-specific gene co-expression networks underpin cognitive performance through distinct biological pathways that vary across the lifespan. Note the convergence of risk and resilience in cell adhesion and nucleic acid regulation across cognitive domains both in children and adults. Also noteworthy is the exclusive association of resilience PGS in stress response pathways in children. Because many cognitive functions rely heavily on synaptic mechanisms, we complemented this analysis with SynGO, a specialized knowledge base focused on synaptic genes and their functions [52]. SynGO provides curated annotations specific to synaptic components and processes, allowing for a more precise evaluation of whether the predictive gene sets are enriched in synapse-related pathways. SynGO analysis revealed an overlap of significant co-expression modules enriched for synaptic pathways across perinatal, juvenile, and adult stages in EUR samples for cognitive flexibility, and across ancestries for the verbal memory domain during perinatal and juvenile stages (Supplementary Figure 10, File SynGO).

**Figure 5:**
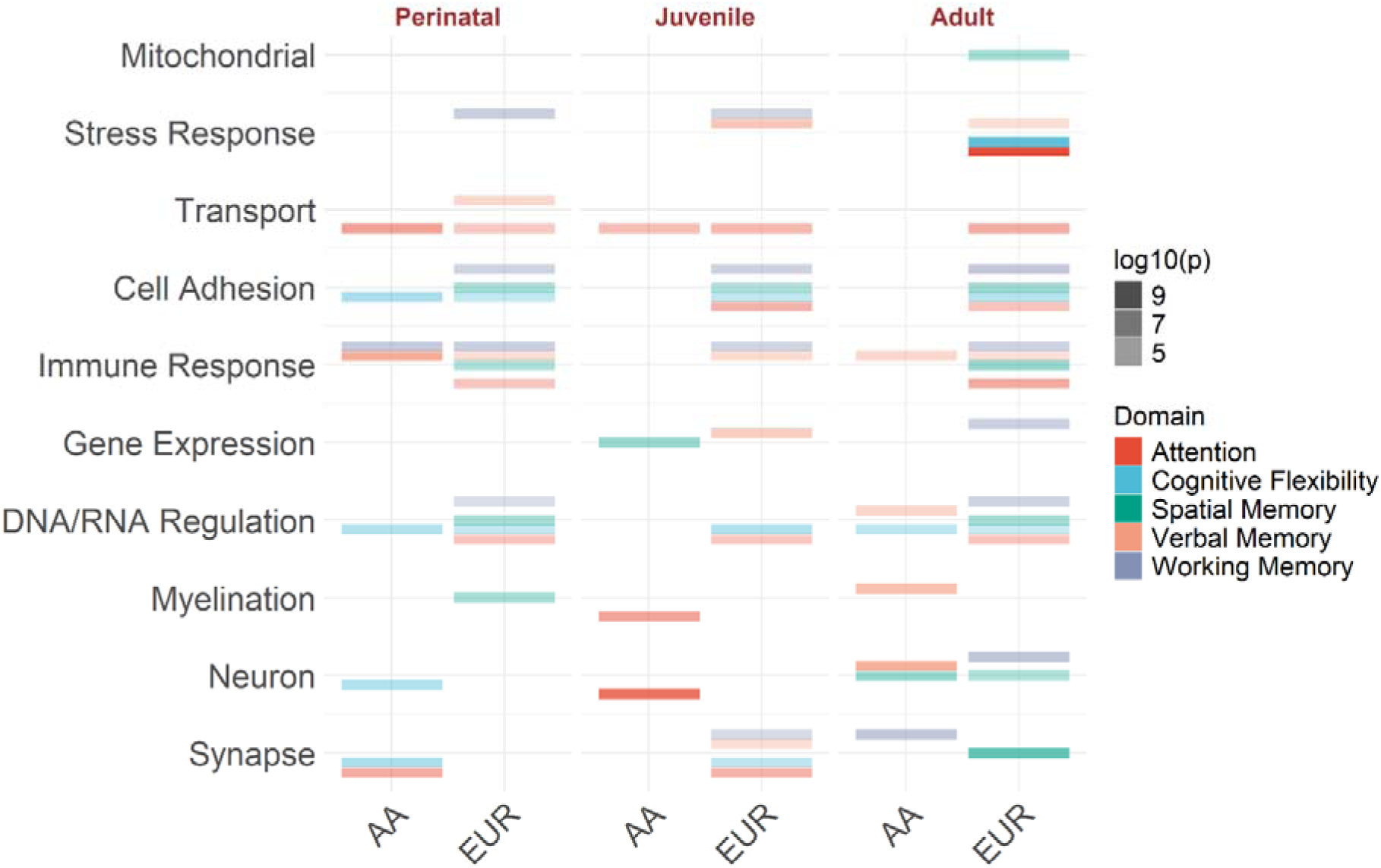
Comparison of GO enrichment for each cognitive domain across three different age stages (perinatal, juvenile and adult) between children/adolescents (ABCD & PNC) in the EUR and AA samples and EUR adults (Bari & NIH). The y-axis represents grouped GO categories while the x-axis represents ethnicities. Tile colors represent different cognitive domains and color intensity increases with a more significant p-value (higher log10(p)).

## Discussion

In this study, we investigated the polygenic effects of SCZ risk variants on cognitive performance across European and African ancestry cohorts of non-psychiatric children, adolescents, and adults; additionally, we evaluated the polygenic effects of SCZ resilience variants in European ancestry individuals. As a positive control, we investigated the polygenic effects of cognitive and general intelligence variants on cognitive performance in the same samples. We aimed to evaluate whether the cognitive impact of SCZ-related genetic architecture is observable across development, whether these associations replicate across individuals of European and African ancestry, and whether the associations differ or converge between risk and resilience variants.

We found robust and replicable negative associations between SCZ risk PGSs and cognitive domains—particularly spatial memory, working memory, verbal memory, and cognitive flexibility— in both European and African individuals, even among individuals whose age preceded the typical onset age of SCZ. These associations were consistent across several cohorts and cognitive domains, underscoring a cross-ancestry convergence of SCZ polygenic risk on key executive and memory-related functions in the absence of the disease and in individuals who most likely will never manifest schizophrenia. Such cross-ancestry replication (e.g., in working memory and cognitive flexibility in ABCD and PNC) highlights the generalizability of SCZ risk-related cognitive underperformance beyond European populations—a critical step toward inclusive psychiatric genetics.

Our findings also show significant opposite associations between SCZ risk and resilience PGSs on cognitive domains, in particular executive function, attention, and working memory. The circumstance that these domains are critically impaired in patients [53–55] ties in with the hypothesized co-segregation of genetic underpinnings of cognitive functioning with SCZ risk [15, 56, 57]. As a positive control, we also found that genetic proxies of cognitive and general intelligence were directly correlated with cognitive performance, especially in children within the domains of working memory, spatial memory, and verbal memory, while showing association in adults only in verbal memory. These results reinforce the idea that the negative associations discovered when considering pathology GWAS scores are not random, but likely reflect genuine biological processes linked to cognitive function and SCZ risk.

Our primary results are consistent with prior research in that deficits in working memory and attention—core features of SCZ—manifest in early childhood as a function of genetic risk (and resilience) [4, 53, 58, 59] in cognitive as well as neuroimaging phenotypes [60]. Additionally, the observed relationship between SCZ polygenic variants and long-term memory deficits corroborates the existing literature [61]. In line with our negative associations with SCZ risk variants and positive associations with intelligence variants, PGSs for SCZ and intelligence have been shown to correlate at the genetic level with a Pearson’s r_g_ = −0.21 [17]. In our case, intelligence-related PGS showed a greater effect size in the association with cognitive abilities in children and adolescents compared to adults, in demographically and geographically distinct cohorts. This pattern contrasts with prior literature suggesting that the heritability of intelligence tends to increase with age [62, 63]. However, it is worth considering that PGS for intelligence explain only part of the overall heritability, and without longitudinal data it is not possible to disentangle age from birth cohort effects.

In the context of prior results on SCZ risk, our findings differ from Germine et al. in 2016 [26], who analyzed the PNC dataset using genetic risk for SCZ derived from the second wave of the Psychiatric Genetic Consortium study. Their results were limited to response times in verbal reasoning and emotion identification tasks. Possible explanations for these differences include our use of PGC3 statistics, our use of the PGScs incorporating Bayesian modeling to better account for ancestry compared to traditional PGSs, including all nominally significant variants, and the inclusion of three additional cohorts to identify the most stable results and extend the investigation across the lifespan. A recent study conducted by Chang et al. [64], which investigated associations between psychiatric or neurodevelopmental PGSs and intra-individual variability in cognitive performances in the ABCD cohort, did not find significant associations between PGS for SCZ and attention or cognitive flexibility. In contrast, our study focused on cross-sectional performance rather than intra-individual variability over time. Other studies, such as He et al. [65], found significant associations between working memory and PGSs for both risk and resilience, while also finding significant results for cognitive flexibility in the Wechsler Memory Scale test. Here, the latter result failed to replicate in the second cohort of adults, even though the same test was used. We did not find any significant interaction with age or sex, despite their well-known role in SCZ incidence and possibly mechanisms [37, 66–68]. The lack of age or sex interactions with the PGSs should be considered in light of the fact that most samples here were of prepuberal age, hence not best-suited to investigate this issue. Nonetheless, our study shows consistency within cognitive domains across different cohorts and suggests that SCZ and cognitive deficits are, in fact, co-heritable, and that these deficits are detectable in an early developmental stage. This notion aligns with the neurodevelopmental hypothesis of SCZ.

To our knowledge, our study is the first to examine parsed polygenic scores for genetic resilience to SCZ to associate age-specific co-expression patterns with cognition. By parsing genetic scores according to age-period relevant co-expression patterns in the brain, polygenic effects previously underrepresented in different cognitive domains, like working memory, spatial memory, verbal memory, and attention in the NIH cohort, emerged as significant. Interestingly, we found cases in which perinatal gene co-expression alone showed explanatory power comparable or superior to that of using all SCZ-associated variants, especially in domains such as working memory in children and attention in adults. These findings were significant across both European and African ancestries, and across childhood and adulthood, pointing to a critical role of early brain development in shaping later cognitive phenotypes linked to SCZ risk. Our findings underscore the importance of early-life molecular contexts in shaping brain function, which we modeled using shifting co-expression patterns across time [33]. Additionally, perinatal co-expression patterns outperformed the explanatory power of adult and juvenile co-expression patterns when pooling all cognitive domains across all cohorts. These results imply a stronger effect on cognitive deficits related to SCZ risk in the perinatal life and highlight the importance of SNP-related effects during the most critical stage of brain development and SCZ susceptibility [69]. Furthermore, even when resilience-associated modules were included (European only), perinatal risk-based modules retained or strengthened their superior explanatory power. This suggests that the impact of early developmental gene expression patterns is a robust feature of SCZ-associated cognitive deficits, as it is observed also when protective genetic factors are accounted for. Interestingly, when parsing cognitive and general intelligence PGSs within the same co-expression architectures, results comparable to SCZ PGS emerge in perinatal co-expression modules, hinting towards shared biological outcomes during the perinatal brain development.

When translating parsed polygenic scores into functional biology through gene ontology analyses, we found that the explanatory co-expression modules for cognition displayed specific functional biology profiles. For example, spatial memory and cognitive flexibility were predicted by parsed scores leveraging variants located in adult co-expression patterns, and cognitive performance was predicted specifically in adult individuals. In contrast, development-related gene enrichment was significant only within juvenile co-expression patterns in younger individuals in association with cognitive flexibility. Interestingly, the interplay between risk and resilience appeared to involve different gene co-expression contexts with separate functional annotations at different stages of life. Overall, the co-expression contexts identified during the perinatal period seemed predominant over contexts identified at later times in the lifespan, as already reported elsewhere [13, 31–33, 70–76]. While most genetic studies have focused on older individuals already experiencing symptoms, our findings suggest that the molecular mechanisms shaping vulnerability or resilience to mental health disorders are already active in early life, when symptoms cannot confound the picture. This perspective emphasizes the potential of early-life interventions for the early identification of risk for SCZ or to enhance resilience in individuals with a genetic predisposition.

We acknowledge the following limitations to our study: first, longitudinal data were available only in the ABCD cohort. As we used cross-sectional data, we were not able to test for replicable cognitive changes or clinical symptoms arising over time at the individual level. Furthermore, while other datasets used in our analysis encompass similar cognitive domains, variations in psychometric tests may influence outcomes, potentially hindering the estimation of the significance of age in cognitive assessments. Moreover, findings regarding genetic resilience are restricted to European ancestry, as the SCZ resilience variants identified by Hess et al. are primarily relevant to European populations. These resilience variants are also not genome-wide significant. In addition, the same SNPs appear in different modules of different age-specific brain region networks, which makes it difficult to establish exclusive relationships within these networks. Lastly, we collapsed our GO enrichment analysis into broader categories due to the extreme diversification of significant pathways, thus obtaining a rather coarse perspective on the biological processes involved.

In summary, our study shows that SCZ polygenic risk is associated with cognitive functioning across development and diverse ancestries and underscores a convergence of SCZ polygenic risk and resilience scores onto the same cognitive domains. Risk and resilience are associated with overlapping biological processes across ancestries, such as transport, cell adhesion, immune response, and DNA regulation in the perinatal age stage. The discovery that SCZ risk-related cognitive deficits are traceable to early life—across ancestries and in the absence of clinical symptoms—raises the question of which preventive strategies can help at-risk youth. Parsing genetic risk by developmental expression patterns also provides a roadmap for identifying windows of vulnerability, and potentially for designing resilience-enhancing interventions. Importantly, genes do not code for SCZ or other disorders; instead, they regulate molecular processes implicated in enabling the brain to interact with its environment [77]. We showed that risk processes are not limited to symptoms characteristic of SCZ onset but additionally implicate the role of risk and resilience in ordinary brain function, well before the onset. These results are strengthened by the specificity of these effects to SCZ and the opposite directions of cognition genetic scores. By integrating gene expression over developmental time, we identify perinatal brain molecular landscapes as particularly relevant for later-life cognition. Together, these findings support a developmentally informed and biologically nuanced view of SCZ risk across ancestries and SCZ resilience, with implications for early identification and intervention.

## Methods

### Population and Datasets

We included genotypes of subjects from four different cohorts ranging in age from 8 to 65 years (mean 14.1), for a total of 16,520 subjects (8,156 females) of European and African ancestry. The first cohort included 6,757 children (3,233 females) between 9 and 12 years of age from the Adolescent Brain and Cognitive Development (ABCD) [78] study. Most ABCD research sites rely on a central Institutional Review Board at the University of California, San Diego for the ethical review and approval of the research protocol, with a few sites obtaining local IRB approval. The second cohort included children and adolescents ranging in age from 8 to 21 years from the Philadelphia Neurodevelopmental Cohort [79] (PNC) study for a total of 7,825 individuals (4,015 females) which were split into two subsets, one comprising 3,070 children (1,416 females) from 8-12 years and one comprising 4,755 adolescents (2,599 females) from 13-21 years. For PNC, all study procedures were approved by the Institutional Review Board at the University of Pennsylvania. In addition, we included 995 healthy adult controls (454 females) from the Apulia region in Italy (BARI), and 943 healthy adult controls (454 females) from the NIH CBDB sibling study [80], ranging in age from 18 to 65 years. For the Bari cohort, the experimental protocol was approved by the institutional ethics committee of the University of Bari Aldo Moro. Written informed consent was obtained after a full understanding of the protocol according to the Declaration of Helsinki. For the NIH cohort, the experimental protocol was approved by the National Institutes of Health research committees. Demographic information for all cohorts included in this study can be found in Table 1. More details on assessment and genotyping procedures can be found elsewhere [70, 71].

**Table 1:**
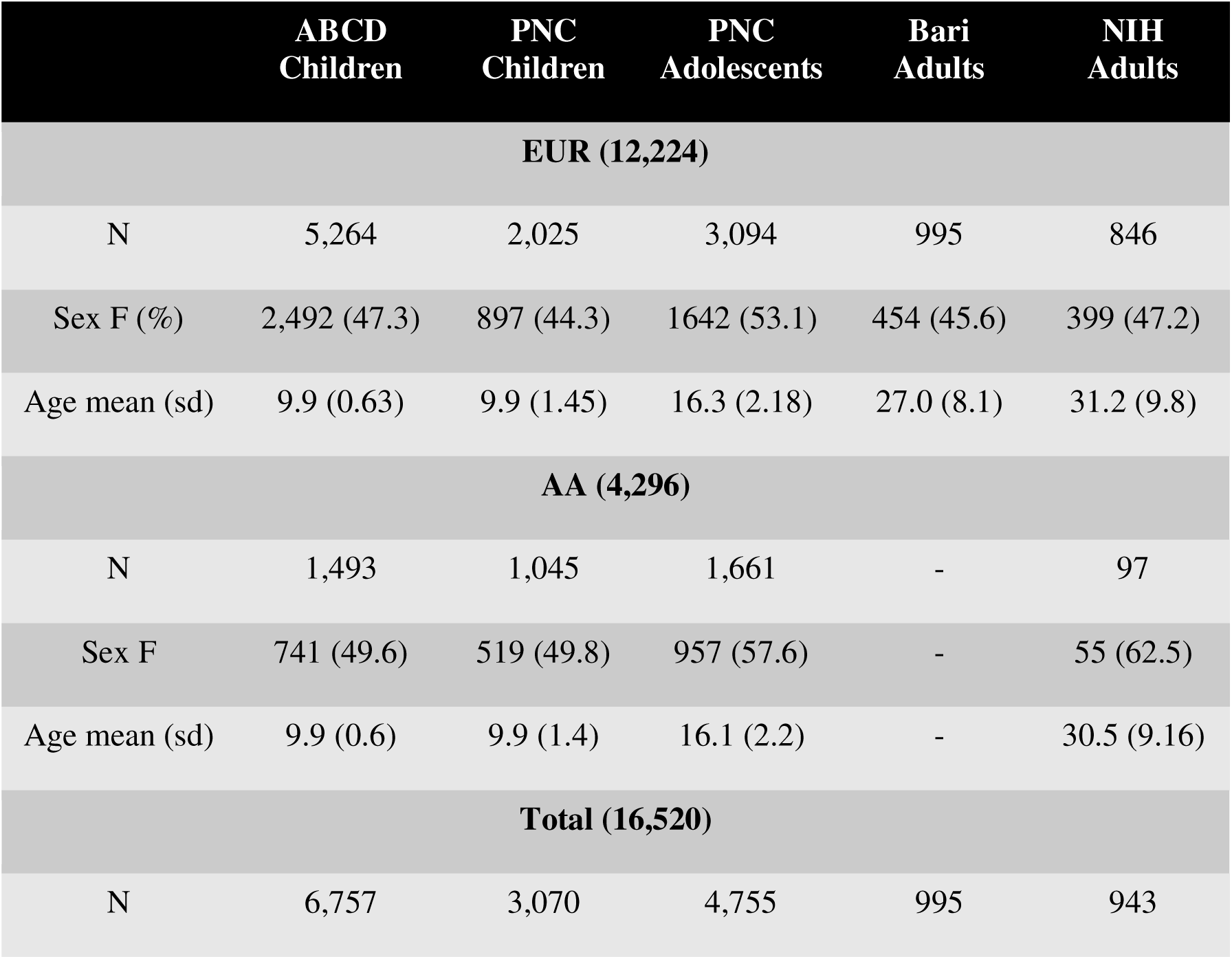
Demographics of all cohorts.

**Table 2.** Effect sizes (t-values) for significant associations between cognitive or general intelligence with cognitive domains in the European subsample.

|  |  | **ABCD**<br>**Children** | **PNC**<br>**Children** | **PNC**<br>**Adolescents** | **Bari**<br>**Adults** | **NIH**<br>**Adults** |
| --- | --- | --- | --- | --- | --- | --- |
| **Attention** | Cognitive Intelligence | 2.4* | 3.3* | 4.4** | n.s. | n.s. |
|  | General Intelligence | n.s. | n.s. | n.s. | n.s. | n.s. |
| **Verbal Memory** | Cognitive Intelligence | 4.6** | 4.9** | 6.8*** | 3.3* | 4.4** |
|  | General Intelligence | 2.1* | 2.6* | 2.4* | n.s. | n.s. |
| **Working Memory** | Cognitive Intelligence | 6.4*** | 5.1** | 7.0*** | n.s. | n.s. |
|  | General Intelligence | 3.7** | 2.7* | 2.9* | n.s. | n.s. |
| **Spatial Memory** | Cognitive Intelligence | 3.7** | 5.6** | 7.4*** | n.s. | n.s. |
|  | General Intelligence | 2.1* | 2.6* | 2.4* | n.s. | n.s. |
| **Cognitive Flexibility** | Cognitive Intelligence | 2.1* | 3.8** | 4.2** | n.s. | n.s. |
|  | General Intelligence | n.s. | n.s. | n.s. | n.s. | n.s. |

ns: $p > 0.05$ , \*: $p \leq 0.05$ , \*\*: $p \leq 0.01$ , \*\*\*: $p \leq 0.001$

All datasets include clinical, cognitive, and genotype data. Inclusion criteria required intact cognitive ability to participate in computerized testing [81].

### Computerized Neurocognitive Battery

All individuals performed neurocognitive tests covering the following cognitive functions: attention, spatial memory, verbal memory, working memory, spatial reasoning and cognitive flexibility. Table 3 illustrates a more detailed description of each test and its cognitive domain. Psychometric data can be found in the supplementary information. In addition, the Wide Range Assessment Test (WRAT) provided an estimate of participants’ IQ. In children, test performances were corrected for parental education status and socioeconomic status.

**Table 3:**
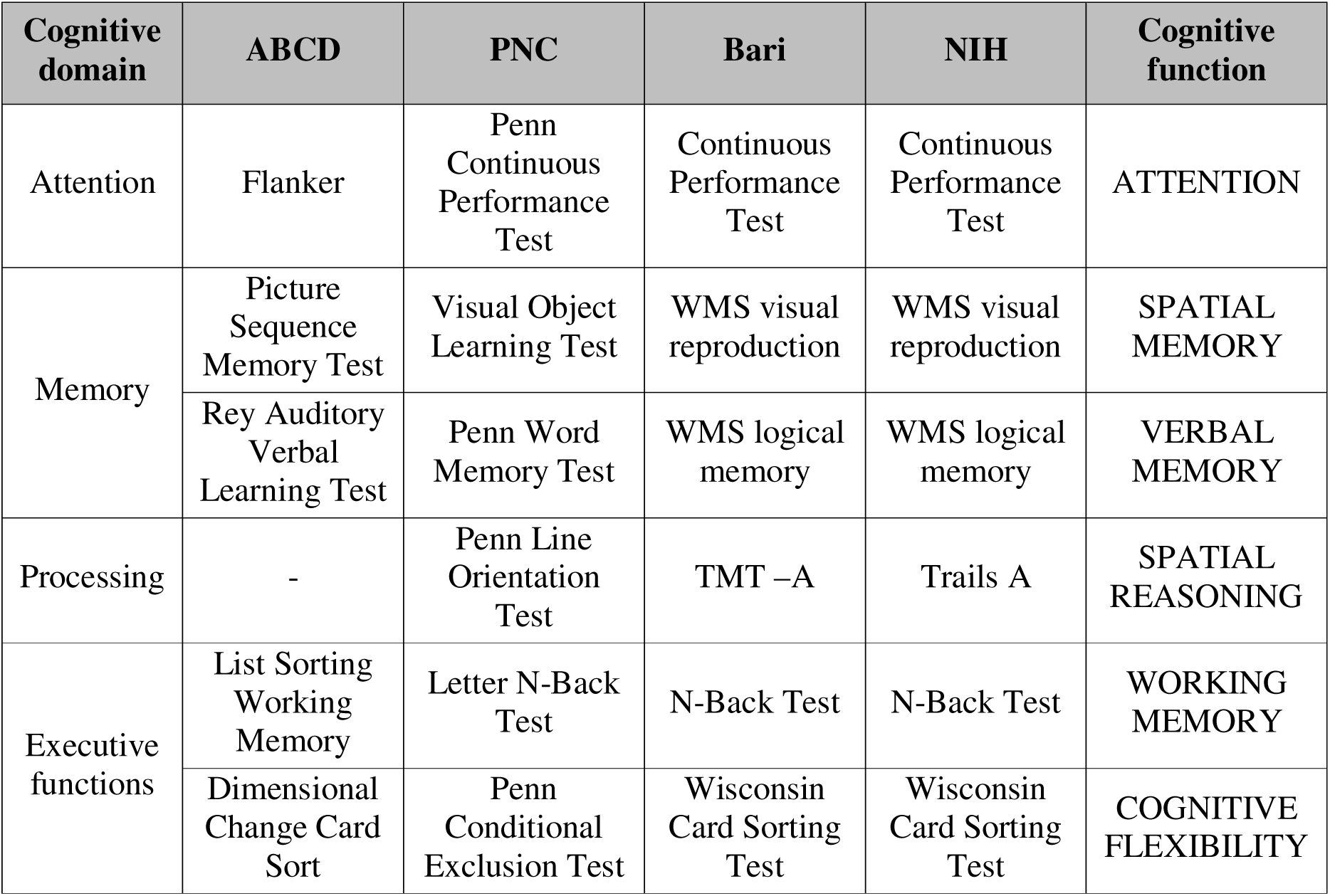
Cognitive tests from all cohorts.

### Quality Control

For all genotyping datasets we applied quality control (QC) following the standard Rapid Imputation and Computational Pipeline (RICOPILI) [82] for genome-wide association studies (GWAS) using PLINK v2 [83] along with other SNP-processing tools on raw genotype data, before imputation as follows: we removed SNPs with a genotype call rate inferior to 95%, individuals with a sample call rate inferior to 95%, and outliers for heterozygosity. Additionally, we removed subjects with non-matching genetic sex corresponding to the reported sex. After subject removal, we removed SNPs with a genotyping quality inferior to 98%, SNPs that are not in Hardy Weinberg Equilibrium (p < 1×10^−6^), and SNPs with a major allele frequency below 1% to exclude rare variants. A more detailed description of each QC step can be found in Supplementary Table 1.

### Ancestry Scores

SCZ resilience variants used in this study for polygenic resilience score calculation have been established based on European ancestry allele frequencies. Therefore, to assess European genetic ancestry, we implemented a cross-validation algorithm for a lasso and elastic-net regularized generalized linear model using HapMap3 [84] as a reference panel, to calculate eigenvalues and assign superpopulations based on overlap of principal components between our sample and the reference sample. Individuals with more than 90% genetic overlap for European ancestry were included in resilience PGS analyses (Supplementary Table 1).

### Imputation

Samples were pre-phased [85] for imputation using the SHAPEIT v1.0.1 [86]: marker strand alignments were checked against the 1000 genomes reference alleles. Any SNPs misaligned with the genomic reference have been flipped prior to phasing to reduce mismatching SNPs during the imputation process. The alternative allele frequency, SNP call rate, and χ2 (reference panel vs. data) are calculated, and allele switches and strand flips are determined, comparing ref/alt from the reference panel with the data. Every chromosome has been imputed separately. Over 500,000 SNPs for each cohort have been used for imputation.

We performed imputation through the Michigan Imputation Server [87], a free imputation service accessible online, which creates chunks of 20 Mb size and executes Minimac4 (performing imputation with pre-phased haplotypes with a MaCH algorithm) for each chunk. As a reference Panel we used the 1000 Genome Phase 3 v5 [88]. The array build was based on GRCh37 / hg19. Phasing for each chunk was done with Eagle v2.4 [89].

After imputation, we retained SNPs with a maximum genotype imputation probability over 0.9 and a minor allele frequency over 1%. We obtained over 4 million SNPs for each dataset.

### Principal Component Analysis

To assess principal component analysis (PCA), we used chip-wide SNP data passing specific quality controls. Hence, we included SNPs common to every different batch (chip array), with an MAF > 1%, a Hardy Weinberg Equilibrium p-value > 10-6, and a missing rate < 2%. We excluded SNPs present in genomic regions known to exhibit extended linkage disequilibrium or structural variation, like the Major Histocompatibility Complex region (Chr6:25-35Mb) and the Chromosome 8 inversion region (Chr8:7-13Mb)[82]. SNPs passing this quality control were pruned to minimize linkage disequilibrium between SNPs (R^2^ < 0.2, 200 SNPs window). With the resulting SNPs, we calculated Eigenvectors and Eigenvalues of the covariance matrix with SNPRelate [90] to identify principal components of the genotype data.

### Polygenic Score Calculation

We chose the continuous shrinkage polygenic risk score (PGScs) approach [91] to include local LD patterns in each dataset and ensure more precise measures of cognitive ability association. PGScs improves upon the classic PGS method by incorporating Bayesian modeling to better account for the complex genetic architecture of traits.

PGScs were calculated with PRSice v2 [92] using the Odds Ratio (OR) obtained from the third wave of the European and African mixed ancestry SCZ GWAS performed by the Psychiatric Genomics Consortium (PGC) [11] and the derived SCZ resilience GWAS [51] as a statistic parameter for allelic scoring. The 1000 genome reference served to estimate the LD during clumping, in which we iteratively picked a SNP (index SNP) and removed variants within a certain genomic window that correlate to the index SNP. We removed SNPs in LD within 500 kb to both ends of the index SNP with an R^2^ threshold of 1 and a p-value threshold of 0.1. One SNP per window is used for PGS calculation. The following formula summarizes the Bayesian model:

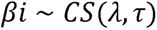

where λ is a global shrinkage parameter, and τ is a local shrinkage parameter that adjusts the degree of shrinkage for each SNP.

Furthermore, we parsed our PGSs based on WGCNA [30] modules within post-mortem brain tissue-derived co-expression networks of different brain regions, e.g., caudate nucleus (CN), dorsolateral prefrontal cortex (DLPFC), and hippocampus, based on different age periods (perinatal, juvenile and adult) to generate age-specific risk profiles and investigate co-expression pattern changes during different stages of life in different brain regions. More detailed methods on RNAseq data and gene co-expression networks derived from the Lieber Institute for Brain Development’s post-mortem brain collection can be found elsewhere [33, 93, 94]. We mapped variants within a range of 100 kbp to each gene inside co-expression modules. We used subsets of SCZ-associated SNPs from genotypes of neurotypical controls to generate as many PGSs as the modules identified in each age- and brain region-specific network. We thus obtained 48 PGSs characterizing perinatal age (fetal to 6 years old), 109 PGSs for juvenile age (6 to 25 years old), and 135 PGSs reflecting adult age (25 to 50 years old).

Finally, we investigated associations of cognitive measures with different neuropsychiatric and neurological disorders, to assess whether our findings would be disease-specific or a result of generic decreased brain functioning. We calculated PGSs for other psychiatric disorders including attention deficit hyperactivity disorder (ADHD), psychotic episodes (single or multiple), major depression disorder (MDD), obsessive compulsive disorder (OCD), bipolar disorder (BD), post-traumatic stress disorder (PTSD) and autism spectrum disorder (ASD), neurodegenerative diseases including Parkinson’s disease (PD) and Alzheimer’s disease (ALZ), non-neurological phenotypes including rheumatoid arthritis (RA), ulcerative colitis (UC), cannabis use disorder (CUD), Crohn’s disease (CD) and height (as a negative control) as well as different measures of intelligence (general intelligence and cognitive functioning) and assessed associations with cognitive performances.

### Linear Models

All statistical analyses employed the R statistical software [95] version 4.2. To compute associations between neurocognitive performance and polygenic risk and resilience scores, we estimated a linear model with the following formula:

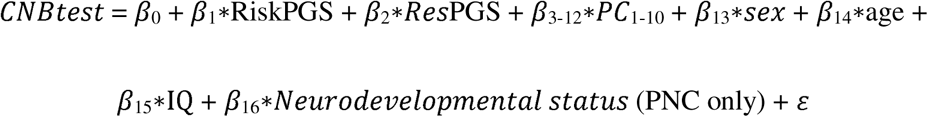

We also investigated possible interactions between risk and resilience with age and sex:

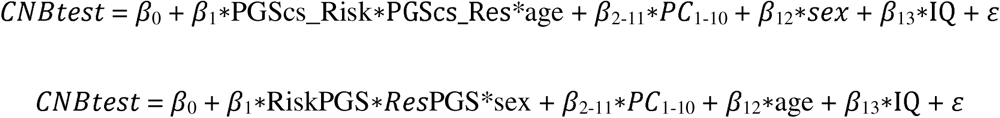

Correlations between each covariate used in this model can be found in Supplementary Figure 9. Each cohort has been analyzed separately to investigate effects relative to each age group. However, we also performed a metanalysis across all cohorts for each cognitive domain using the metafor [96] and metap R package [97] on each coefficients p-value.

Performance measures from six different cognitive domains (attention, working memory, spatial memory, spatial memory, cognitive flexibility, verbal reasoning, and spatial reasoning) were used as dependent variables. Standardized SCZ risk (PGScs_Risk) and resilience (PGScs_Res) polygenic scores were used as independent variables. Further, we included the first 10 PCs, biological sex, intelligence quotient, and subjects’ age. We removed outliers for performances and polygenic scores by calculating z-scores and excluding values over three standard deviations from zero. The resulting p-values for every linear model were corrected for the total number of linear models with the Bonferroni correction method for number of traits (risk and resilience) and number of cognitive domains (six), hence a total of 12 comparisons. We used the T-values from the R linear model as a measure of the effect sizes. T-values in this function are calculated by dividing the b coefficient of a linear model by its estimated standard error. Positive numbers represent positive associations, and negative signs represent negative associations. Delta t-values were obtained by subtracting resilience from risk t estimates, resulting in positive delta t-values representing divergent effects, i.e., positive resilience and negative risk scores. To investigate the divergence between risk and resilience PGSs, we calculated the difference between their effect sizes (Δ t-values) and assessed the differences of their β estimates with a linear hypothesis test (LHT), using the linearHypothesis() function in the ‘car’ package. LHT p-values have been FDR corrected.

Finally, we established a machine learning pipeline to assess whether a combination of parsed co-expression modules for specific age periods across three brain regions (WGCNA) could explain more variance in cognitive performances across different age groups. As such, we divided each cohort into two subsets (k = 2-fold) and used one subset as a training sample to extract significant PGSs parsed for age-relevant co-expression modules through a linear model and predict each cognitive test on the testing sample. We performed 100 iterations and calculated the Akaike Information Criterion (AIC) of the full PGScs and parsed PGScs models for each iteration, as well as Pearson’s correlation R between observed and predicted cognitive performance by full and parsed PGScs. To establish the best performance, we calculated the AIC difference (Δ_AIC_) between both models (PGS_whole_ Δ_AIC_ – PGS_parsed_ Δ_AIC_) and assessed significance through a Wilcoxon signed-rank test. For WGCNA, we evaluated three different models: Co-expression modules parsed in adult, juvenile, and perinatal lifespan, including DLPFC, HP, and CN (the latter was available for adult and juvenile samples only). We also calculated the variance explained by the parsed models on the full PGScs model’s residuals and vice versa to control for the overlapping variance explained by both models.

Lastly, we asked which biological functions the predictive gene sets tapped into. To this end, we extracted all the significant (p < 0.05) parsed PGSs for each age stage (perinatal, juvenile, and adult) from each of the 100 iterations and chose the top 5% most significant scores (each score representing a co-expression module) for perinatal, juvenile and adults across all iterations that were also significant in at least 50% of all iterations. We then pooled together the genes included in the top 5% significant co-expression modules for each age period and cognitive domain and performed gene ontology (GO) enrichment analysis with the ClusterProfiler [98] R package. We tested FDR-corrected enrichment p-values with a Fisher’s combined probability test in the adult and non-adult individuals of our four cohorts, respectively, and chose enriched ontologies with a significance of Fisher’s p < 0.001. In addition to GO enrichment, we performed enrichment analysis using SynGO [52], a curated knowledge base dedicated to synaptic genes and functions. SynGO contains experimentally validated annotations for synaptic components and processes, allowing for a more targeted assessment of synapse-specific biological pathways compared to general GO terms. We used the SynGO web platform (https://syngoportal.org) to test whether predictive gene sets identified in each age stage were significantly enriched for synaptic annotations, using the default settings and a false discovery rate (FDR) of 5% for multiple-testing correction.

## Supporting information

Supplementary Psychometric Data

## Acknowledgements

We would like to thank Exprivia S.p.A. for supporting G.C.K with a scholarship throughout his PhD. We are grateful for the contributions of Dr. Qiang Chen for his help in establishing scripts and pipelines for genotype data preparation and Dr. Nora Penzel for sharing her knowledge of RStudio plotting and tidying scripts. We acknowledge the contribution of Prof. Giuseppe Blasi to the collection of genetic and cognitive data from Bari and Dr. Roberta Passiatore to the curation of cognitive and demographic data from NIH. The Philadelphia Neurodevelopmental Cohort (PNC) is a research initiative funded by the NIMH through the American Reinvestment and Recovery Act of 2009 (www.med.upenn.edu/bbl/philadelphianeurodevelopmentalcohort). The initiative focuses on characterizing brain and behavior interaction with genetics. This is a collaborative research effort between the Brain Behavior Laboratory at the University of Pennsylvania and the Center for Applied Genomics at the Children’s Hospital of Philadelphia (CHOP), led by Raquel E. Gur, MD, Ph.D. at the University of Pennsylvania and Hakon Hakonarson, MD, Ph.D., at the CHOP. Permission to use the PNC was provided to G.P. under project ID: 20998. Data from the ABCD Study®, held in the NIMH Data Archive (NDA), were obtained under approved request No 12036. The ABCD Study® is supported by the NIH and additional federal partners under award numbers U01DA041048, U01DA050989, U01DA051016, U01DA041022, U01DA051018, U01DA051037, U01DA050987, U01DA041174, U01DA041106, U01DA041117, U01DA041028, U01DA041134, U01DA050988, U01DA051039, U01DA041156, U01DA041025, U01DA041120, U01DA051038, U01DA041148, U01DA041093, U01DA041089, U24DA041123, and U24DA041147. A full list of supporters is available at https://abcdstudy.org/federal-partners.html. A listing of participating sites and a complete listing of the study investigators can be found at abcdstudy.org/consortium_members/. ABCD consortium investigators designed and implemented the study and/or provided data but did not necessarily participate in the analysis or writing of this manuscript. The ABCD data repository grows and changes over time. The ABCD data used in this report came from NIMH Data Archive DOI: https://doi.org/10.15154/1528793. The collection of the genetic and cognitive data for the NIH cohort was supported by direct funding from the Intramural Research Program of the NIMH to the Clinical Brain Disorders Branch (PI: D.R.W., protocol 95-M-0150).

## Data availability

Permission to use the PNC was provided to G.P. under project ID: 20998 and genetic and cognitive data are available upon approved request at the following link: www.med.upenn.edu/bbl/philadelphianeurodevelopmentalcohort. Data from the ABCD Study®, held in the NIMH Data Archive, were obtained under approved request No 12036 and can be accessed upon request at the following link: https://abcdstudy.org/. ABCD data used in this report came from NIMH Data Archive https://doi.org/10.15154/1528793. Genetic and cognitive data from Bari cannot be shared at the individual level in raw format because of ethical restrictions based on the protocol approved by the institutional ethics committee of the UNIBA to protect the privacy of the participants. The individual deidentified raw data from NIH analyzed during this study are available from the corresponding author G.P. upon reasonable request. The GWAS summary statistics used in this study are available at : <u>pgc.unc.edu/for-researchers/download-results/</u>, https://www.ebi.ac.uk/gwas/home and portals.broadinstitute.org/collaboration/giant/index.php/. Summary statistics for cognitive intelligence are available in the supplementary data from <u>10.1038/s41467-018-04362-x</u> and for general intelligence in the supplementary data from https://doi.org/10.1038/s41588-018-0152-6.

## Code Availability

The scripts used for the analyses conducted in this study are available online upon request under the following link: https://doi.org/10.5281/zenodo.14655751.

## References

1. van Os, J., G. Kenis, and B.P.F. Rutten, The environment and schizophrenia. Nature, 2010. 468(7321): p. 203–212.

2. Zhou, J., et al., Working memory deficits in children with schizophrenia and its mechanism, susceptibility genes, and improvement: A literature review. Frontiers in psychiatry, 2022. 13.

3. Tripathi, A., S.K. Kar, and R. Shukla, Cognitive Deficits in Schizophrenia: Understanding the Biological Correlates and Remediation Strategies. 2018.

4. Forbes, N., Working memory in schizophrenia: a meta-analysis | Psychological Medicine | Cambridge Core. Psychological Medicine, 2009.

5. Cameron, A., et al., Working memory correlates of three symptom clusters in schizophrenia. Psychiatry research, 2002. 110(1).

6. Bora, E., Differences in cognitive impairment between schizophrenia and bipolar disorder: Considering the role of heterogeneity. Psychiatry and clinical neurosciences, 2016. 70(10).

7. Tandon, R., et al., Definition and description of schizophrenia in the DSM-5. Schizophrenia research, 2013. 150(1).

8. Häfner, H., et al., IRAOS: an instrument for the assessment of onset and early course of schizophrenia. Schizophrenia research, 1992. 6(3).

9. Green, M., et al., Neurocognitive deficits and functional outcome in schizophrenia: are we measuring the “right stuff“? Schizophrenia bulletin, 2000. 26(1).

10. Sullivan, P., M. Daly, and M. O’Donovan., Genetic architectures of psychiatric disorders: the emerging picture and its implications. Nature reviews. Genetics, 2012. 13(8).

11. Trubetskoy, V., et al., Mapping genomic loci implicates genes and synaptic biology in schizophrenia. Nature, 2022. 604(7906): p. 502–508.

12. Ripke, S. and S.W.G.o.t.P.G. Consortium, Biological insights from 108 schizophrenia-associated genetic loci. Nature, 2014. 511(7510).

13. Richards, A.L., et al., The Relationship Between Polygenic Risk Scores and Cognition in Schizophrenia. Schizophrenia Bulletin, 2023. 46(2): p. 336–344.

14. Smeland, O.B., et al., Identification of Genetic Loci Jointly Influencing Schizophrenia Risk and the Cognitive Traits of Verbal-Numerical Reasoning, Reaction Time, and General Cognitive Function. JAMA Psychiatry, 2023. 74(10): p. 1065–1075.

15. Snitz, B.E., A.W. Macdonald, and C.S. Carter, Cognitive deficits in unaffected first-degree relatives of schizophrenia patients: a meta-analytic review of putative endophenotypes. Schizophrenia bulletin, 2006. 32(1).

16. Harrison, P.J. and D.R. Weinberger, Schizophrenia genes, gene expression, and neuropathology: on the matter of their convergence. Molecular Psychiatry, 2004. 10(1): p. 40–68.

17. Davies, G., et al., Study of 300,486 individuals identifies 148 independent genetic loci influencing general cognitive function. Nature Communications, 2018. 9(1): p. 1–16.

18. Savage, J.E., et al., Genome-wide association meta-analysis in 269,867 individuals identifies new genetic and functional links to intelligence. Nature Genetics, 2018. 50(7): p. 912–919.

19. Blokland, G.A.M., et al., Heritability of Neuropsychological Measures in Schizophrenia and Nonpsychiatric Populations: A Systematic Review and Meta-analysis. Schizophrenia Bulletin, 2017. 43(4): p. 788–800.

20. Bora, E., et al., Cognitive deficits in youth with familial and clinical high risk to psychosis: a systematic review and meta-analysis. Acta psychiatrica Scandinavica, 2014. 130(1).

21. Toulopoulou, T., et al., Polygenic risk score increases schizophrenia liability through cognition-relevant pathways. Brain, 2019. 142(2): p. 471–485.

22. Engen, M.J., et al., Polygenic scores for schizophrenia and general cognitive ability: associations with six cognitive domains, premorbid intelligence, and cognitive composite score in individuals with a psychotic disorder and in healthy controls. Translational Psychiatry, 2020. 10(1): p. 1–9.

23. Fatemi, S. and T. Folsom, The neurodevelopmental hypothesis of schizophrenia, revisited. Schizophrenia bulletin, 2009. 35(3).

24. Weinberger, D.R., Implications of normal brain development for the pathogenesis of schizophrenia. Archives of general psychiatry, 1987. 44(7).

25. Gottesman, I.I. and J. Shields, A polygenic theory of schizophrenia. Proceedings of the National Academy of Sciences of the United States of America, 1967. 58(1): p. 199.

26. Germine, L., et al., Association between polygenic risk for schizophrenia, neurocognition and social cognition across development. Translational psychiatry, 2016. 6(10).

27. Weinberger, D., The neurodevelopmental origins of schizophrenia in the penumbra of genomic medicine. 2017, World Psychiatry.

28. Nasrallah, H.A. and D.R. Weinberger, The neurology of schizophrenia. 1986.

29. Birnbaum, R. and D.R. Weinberger, The Genesis of Schizophrenia: An Origin Story. 10.1176/appi.ajp.20240305, 2024.

30. Li, J., et al., Application of Weighted Gene Co-expression Network Analysis for Data from Paired Design. Scientific Reports, 2018. 8(1): p. 1–8.

31. Walker, R., Genetic Control of Expression and Splicing in Developing Human Brain Informs Disease Mechanisms: Cell. 2019.

32. Werling, D., Whole-Genome and RNA Sequencing Reveal Variation and Transcriptomic Coordination in the Developing Human Prefrontal Cortex: Cell Reports. 2020.

33. Pergola, G., Consensus molecular environment of schizophrenia risk genes in coexpression networks shifting across age and brain regions. 2023.

34. Gandal, M.J., et al., Transcriptome-wide isoform-level dysregulation in ASD, schizophrenia, and bipolar disorder. 2018.

35. Nour, M. and O. Howes, Interpreting the neurodevelopmental hypothesis of schizophrenia in the context of normal brain development and ageing. Proceedings of the National Academy of Sciences of the United States of America, 2015. 112(21).

36. Clifton, N.E., et al., Dynamic expression of genes associated with schizophrenia and bipolar disorder across development. Translational Psychiatry, 2019. 9(1): p. 1–9.

37. Passiatore, R., et al., Changes in patterns of age-related network connectivity are associated with risk for schizophrenia. 2023.

38. Susan, K., Age-dependent patterns of schizophrenia genetic risk affect cognition.

39. Xilin Jiang, C.H., Gil McVean, The impact of age on genetic risk for common diseases | PLOS Genetics. PLOS Genetics, 2021.

40. Meer, D.v.d., et al., Clustering Schizophrenia Genes by Their Temporal Expression Patterns Aids Functional Interpretation. Schizophrenia Bulletin, 2023. 50(2): p. 327.

41. Colantuoni, C., et al., Temporal dynamics and genetic control of transcription in the human prefrontal cortex. Nature, 2011. 478(7370).

42. Kang, H.J., et al., Spatio-temporal transcriptome of the human brain. Nature, 2011. 478(7370).

43. Pergola, G., et al., Lessons Learned From Parsing Genetic Risk for Schizophrenia Into Biological Pathways. Biol Psychiatry, 2022.

44. Robinson, N., et al., Impact of Early-Life Factors on Risk for Schizophrenia and Bipolar Disorder. Schizophrenia Bulletin, 2023. 49(3): p. 768–777.

45. Sportelli, L., et al., Dopamine signaling enriched striatal gene set predicts striatal dopamine synthesis and physiological activity in vivo. Nature Communications, 2024. 15(1): p. 1–19.

46. Martin, A.R., et al., Clinical use of current polygenic risk scores may exacerbate health disparities. Nature Genetics, 2019. 51(4): p. 584–591.

47. Bigdeli, T.B., et al., Contributions of common genetic variants to risk of schizophrenia among individuals of African and Latino ancestry. Molecular Psychiatry, 2019. 25(10): p. 2455–2467.

48. RE, P., et al., Genome-wide Association Studies in Ancestrally Diverse Populations: Opportunities, Methods, Pitfalls, and Recommendations. Cell, 2019. 179(3).

49. Bigdeli, T.B., et al., Biological Insights from Schizophrenia-associated Loci in Ancestral Populations. 2024.

50. Benjamin, K.J.M., et al., Analysis of gene expression in the postmortem brain of neurotypical Black Americans reveals contributions of genetic ancestry. Nature Neuroscience, 2024. 27(6): p. 1064–1074.

51. Hess, J.L., et al., A polygenic resilience score moderates the genetic risk for schizophrenia. Molecular Psychiatry, 2019. 26(3): p. 800–815.

52. F, K., et al., SynGO: An Evidence-Based, Expert-Curated Knowledge Base for the Synapse. Neuron, 2019. 103(2).

53. Luck, S. and J. Gold, Attention in Schizophrenia. Current topics in behavioral neurosciences, 2023. 63.

54. Forbes, N., Working memory in schizophrenia: a meta-analysis | Psychological Medicine | Cambridge Core. 2009.

55. Braun, U., et al., Brain network dynamics during working memory are modulated by dopamine and diminished in schizophrenia. Nature Communications, 2021. 12(1): p. 1–11.

56. Callicott, J.H., et al., Abnormal fMRI Response of the Dorsolateral Prefrontal Cortex in Cognitively Intact Siblings of Patients With Schizophrenia. 10.1176/appi.ajp.160.4.709, 2003.

57. Krista M., Wisner, B.E., James M. Gold, Daniel R. Weinberger, Dwight Dickinson, A closer look at siblings of patients with schizophrenia: The association of depression history and sex with cognitive phenotypes. 2011, Schizophrenia Research.

58. Carter, J.e.a., Attention deficits in schizophrenia — Preliminary evidence of dissociable transient and sustained deficits Author links open overlay panel. 2010, Schizophrenia Research.

59. Gold, J. and S. Luck, Working Memory in People with Schizophrenia | SpringerLink. 2022: SpringerLink.

60. Gur, R., Functional magnetic resonance imaging in schizophrenia. 2010, Dialogues Clin Neurosci.

61. Van de Ven, V., Reduced intrinsic visual cortical connectivity is associated with impaired perceptual closure in schizophrenia. 2017.

62. Bouchard, T., The Wilson Effect: The Increase in Heritability of IQ With Age | Twin Research and Human Genetics | Cambridge Core. Twin Research and Human Genetics, 2013.

63. Haworth, C., et al., The heritability of general cognitive ability increases linearly from childhood to young adulthood. Molecular psychiatry, 2009. 15(11): p. 1112.

64. Chang, S.E., et al., Attention-mediated genetic influences on psychotic symptomatology in adolescence. Nature Mental Health, 2024: p. 1–14.

65. He, Q., et al., Influence of polygenic risk scores for schizophrenia and resilience on the cognition of individuals at-risk for psychosis. Translational Psychiatry, 2021. 11(1): p. 1–9.

66. Ursini, G., et al., Prioritization of potential causative genes for schizophrenia in placenta. Nature Communications, 2023. 14(1): p. 1–17.

67. Geraci, F., et al., Sex dimorphism controls dysbindin-related cognitive dysfunctions in mice and humans with the contribution of COMT. Molecular Psychiatry, 2024. 29(9): p. 2666–2677.

68. Li, X., et al., A glimpse of gender differences in schizophrenia. General Psychiatry, 2022. 35(4).

69. Aguilar-Valles, A., B. Rodrigue, and E. Matta-Camacho, Frontiers | Maternal Immune Activation and the Development of Dopaminergic Neurotransmission of the Offspring: Relevance for Schizophrenia and Other Psychoses. 2020.

70. Pergola, G., et al., DRD2 co-expression network and a related polygenic index predict imaging, behavioral and clinical phenotypes linked to schizophrenia. Translational Psychiatry, 2017. 7(1).

71. Pergola, G., et al., Prefrontal Coexpression of Schizophrenia Risk Genes Is Associated With Treatment Response in Patients. Biological psychiatry, 2019. 86(1).

72. Li, M., et al., A human-specific AS3MT isoform and BORCS7 are molecular risk factors in the 10q24.32 schizophrenia-associated locus. Nat Med, 2016. 22(6): p. 649–56.

73. Fromer, M., et al., Gene expression elucidates functional impact of polygenic risk for schizophrenia. Nature neuroscience, 2016. 19(11).

74. Gandal, M.J., et al., Shared molecular neuropathology across major psychiatric disorders parallels polygenic overlap. Science, 2018. 359(6376): p. 693-697.

75. Hartl, C.L., et al., Coexpression network architecture reveals the brain-wide and multiregional basis of disease susceptibility. Nature neuroscience, 2021. 24(9).

76. Radulescu, E., et al., Identification and prioritization of gene sets associated with schizophrenia risk by co-expression network analysis in human brain. Molecular Psychiatry, 2018. 25(4): p. 791–804.

77. Weinberger, D.R., Thinking About Schizophrenia in an Era of Genomic Medicine. Am J Psychiatry, 2019. 176(1): p. 12–20.

78. Jernigan, T., B. SA, and D. GJ, The Adolescent Brain Cognitive Development Study. Journal of research on adolescence : the official journal of the Society for Research on Adolescence, 2018. 28(1).

79. Satterthwaite, T., et al., The Philadelphia Neurodevelopmental Cohort: A publicly available resource for the study of normal and abnormal brain development in youth. NeuroImage, 2016. 124(Pt B).

80. Dickinson, D., et al., Cognitive Factor Structure and Invariance in People With Schizophrenia, Their Unaffected Siblings, and Controls. Schizophrenia Bulletin, 2011. 37(6): p. 1157–1167.

81. Calkins, M., et al., The Philadelphia Neurodevelopmental Cohort: constructing a deep phenotyping collaborative. Journal of child psychology and psychiatry, and allied disciplines, 2015. 56(12).

82. Lam, M., et al., RICOPILI: Rapid Imputation for COnsortias PIpeLIne. Bioinformatics (Oxford, England), 2020. 36(3).

83. Purcell, S., et al., PLINK: a tool set for whole-genome association and population-based linkage analyses. American journal of human genetics, 2007. 81(3).

84. Duan, S., et al., FstSNP-HapMap3: a database of SNPs with high population differentiation for HapMap3. Bioinformation, 2008. 3(3).

85. Howie, B., et al., Fast and accurate genotype imputation in genome-wide association studies through pre-phasing. Nature genetics, 2012. 44(8).

86. Delaneau, O., J. Marchini, and J.-F. Zagury, A linear complexity phasing method for thousands of genomes. Nature Methods, 2011. 9(2): p. 179–181.

87. Das, S., et al., Next-generation genotype imputation service and methods. Nature genetics, 2016. 48(10).

88. Clarke, L., et al., The international Genome sample resource (IGSR): A worldwide collection of genome variation incorporating the 1000 Genomes Project data. Nucleic acids research, 2017. 45(D1).

89. Loh, P.-R., et al., Reference-based phasing using the Haplotype Reference Consortium panel. Nature Genetics, 2016. 48(11): p. 1443–1448.

90. Zheng, X., et al., A high-performance computing toolset for relatedness and principal component analysis of SNP data. Bioinformatics (Oxford, England), 2012. 28(24).

91. Tian, G., et al., Polygenic prediction via Bayesian regression and continuous shrinkage priors. Nature Communications, 2019. 10(1): p. 1–10.

92. Choi, S. and P. O’Reilly, PRSice-2: Polygenic Risk Score software for biobank-scale data. GigaScience, 2019. 8(7).

93. Collado-Torres, L., Regional Heterogeneity in Gene Expression, Regulation, and Coherence in the Frontal Cortex and Hippocampus across Development and Schizophrenia. 2019.

94. Jaffe, A.E., et al., qSVA framework for RNA quality correction in differential expression analysis. 2017.

95. R Core Team (2020) R A Language and Environment for Statistical Computing. R Foundation for Statistical Computing, Vienna, Austria. - References - Scientific Research Publishing. 2023; Available from: https://scirp.org/reference/referencespapers.aspx?referenceid=3064798.

96. Viechtbauer, W., Conducting meta-analyses in R with the metafor package. Journal of Statistical Software, 2010.

97. Dewey, M., metap: Meta-Analysis of Significance Values. 2024. p. https://CRAN.R-project.org/package=metap.

98. Wu T, H.E., Xu S, Chen M, Guo P, Dai Z, Feng T, Zhou L, Tang W, Zhan L, Fu x, Liu S, Bo X, Yu G. clusterProfiler 4.0: A universal enrichment tool for interpreting omics data. 2021; Available from: http://bioconductor.org/packages/clusterProfiler/.

