## Supplementary Figures and Tables for "Genetic Risk and Resilience for Schizophrenia Stratified by Perinatal Gene Expression Predict Adult Cognitive Performance"

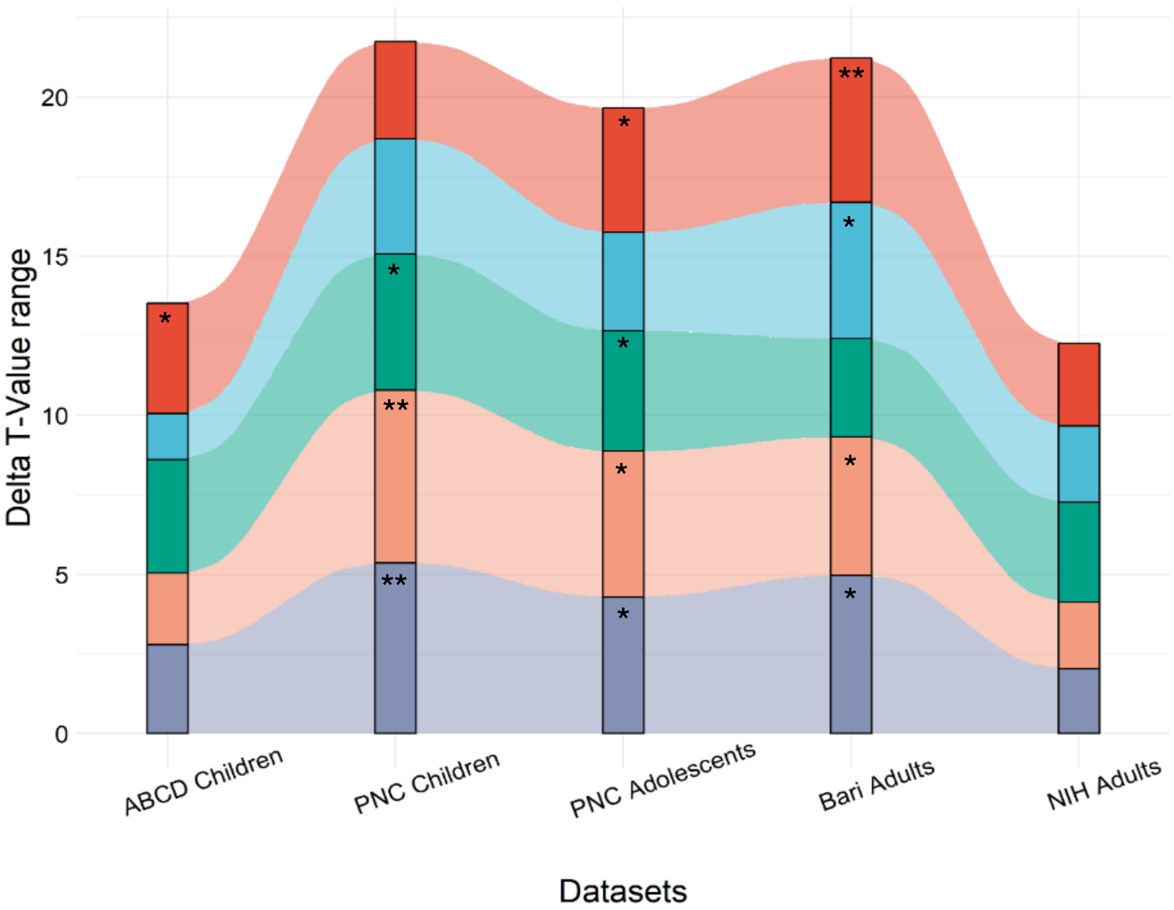


Supplementary figure 1: Risk and resilience Delta PGS effects in the ABCD, PNC, Bari and NIH cohorts on all cognitive domains using the PGC wave 2 summary statistics for both SCZ risk and resilience variants. Delta for T values for each score are shown on the y-axis. Associations are statistically significant in attention, cognitive flexibility, spatial memory, verbal memory and working memory performance. Asterisks signal the significance of linear hypothesis tests between risk and resilience estimates (*: $p < 0.05$, **: $p <= 0.01$, ***: $p <= 0.001$). Connections between bar plots highlight the changes in effect sizes between different cohorts ordered by median age from left to right.

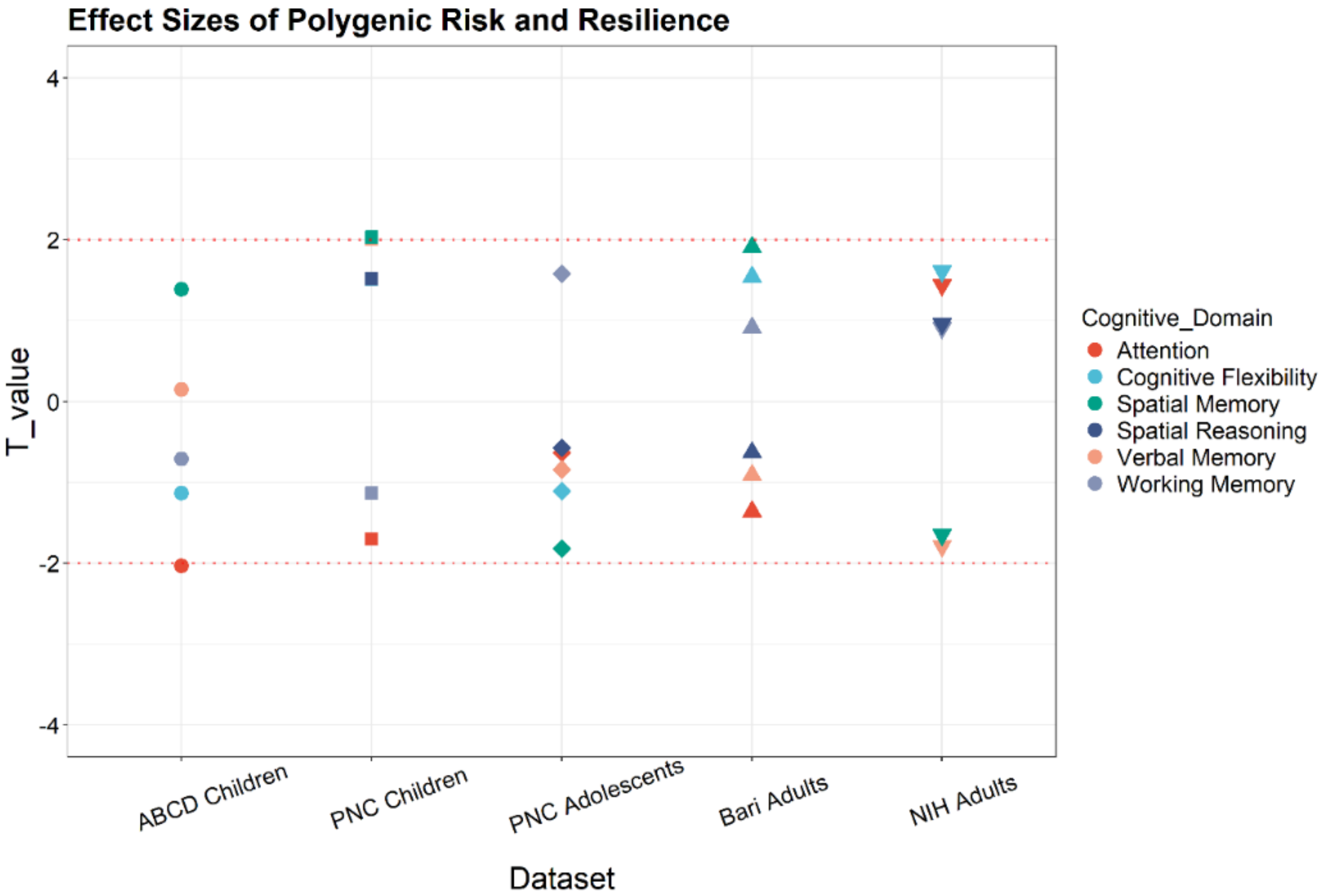


Supplementary figure 2: Interaction between risk and resilience with cognitive domains

*Effect sizes of polygenic risk and resilience interaction on cognitive domains across all cohorts. There is no significant effect of risk-resilience interaction on any cognitive domain (all p>0.05).*

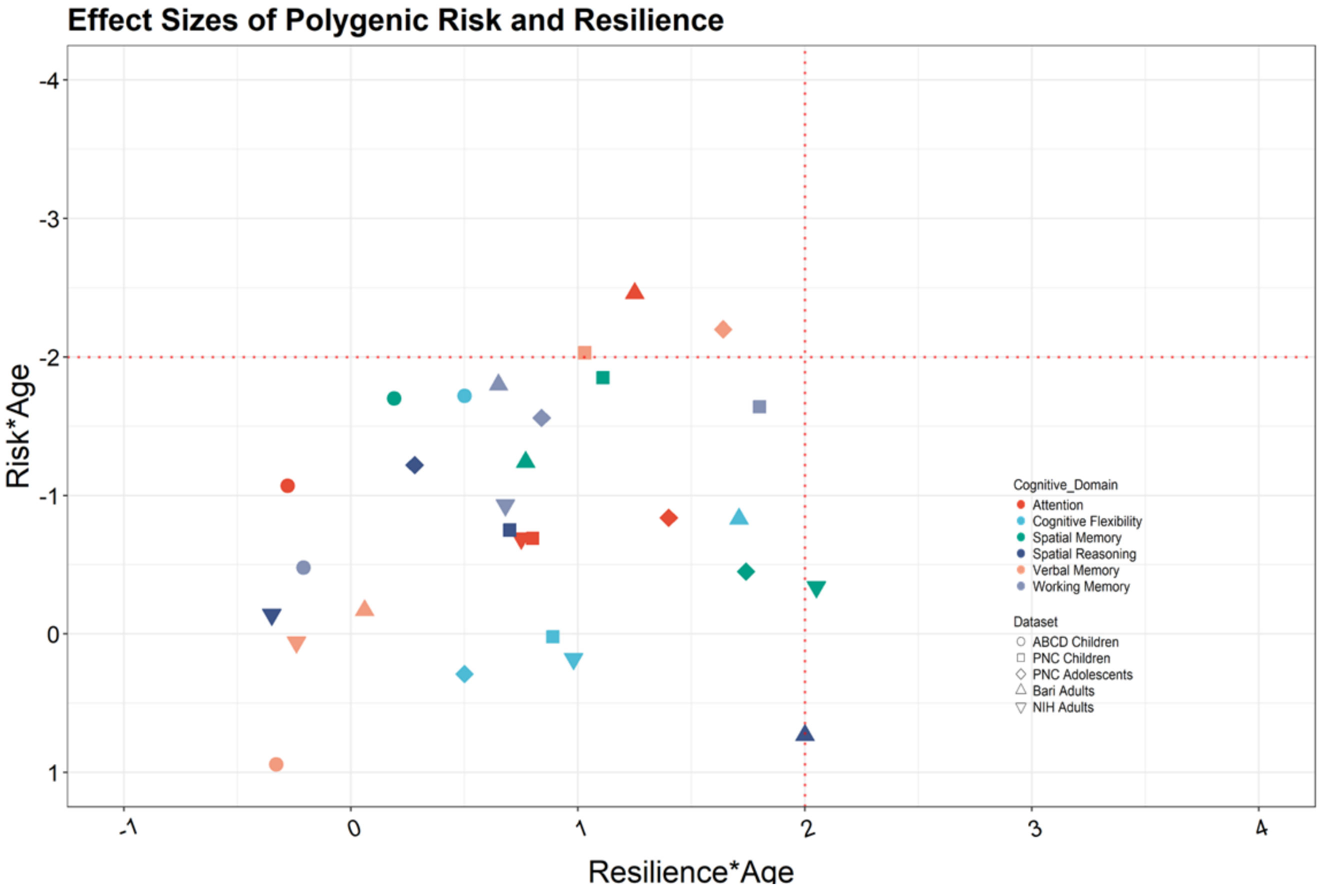


Supplementary figure 3: Interaction between risk, resilience and age with cognitive domains

*Effect sizes of age interaction with polygenic risk (y-axis) and resilience (x-axis) on cognitive domains across all cohorts. There is a significant effect of age-risk in the adult cohort from Bari in attention performance ($t = -2.46$, $p < .01$), and within the children and adolescent cohort from PNC regarding verbal memory performance ($t = -2.03$, $p < 0.03$, $t = -2.2$, $p < .02$). No risk-resilience or age-resilience interaction on any cognitive domain remain significant after multiple comparison correction (all $p>0.05$).*

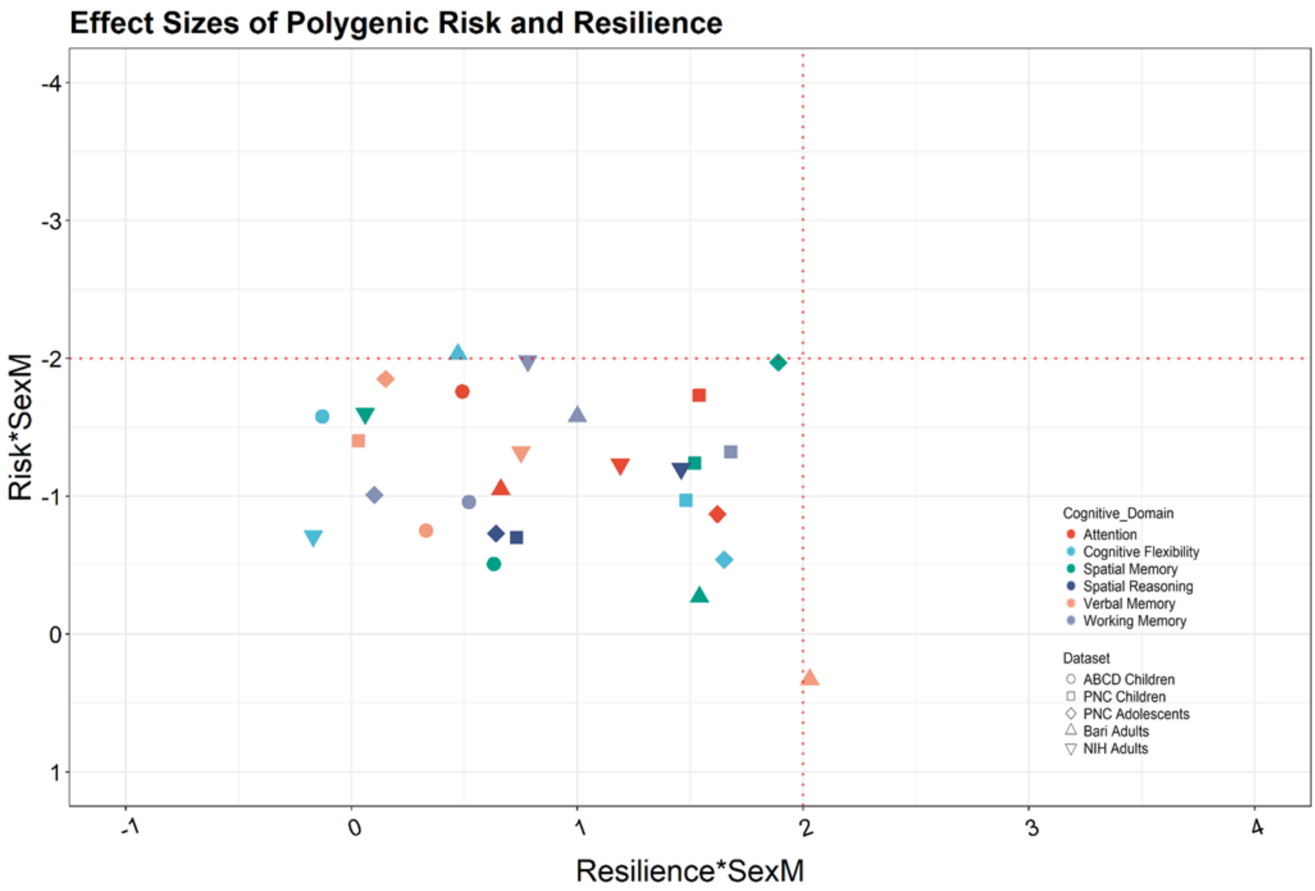


Supplementary figure 4: Interaction between risk resilience and sex with cognitive domains

*Effect sizes of biological sex interaction with polygenic risk (y-axis) and resilience (x-axis) on cognitive domains across all cohorts. There is no significant effect of sex-risk or sex-resilience interaction on any cognitive domain after multiple comparison correction (all p>0.05).*

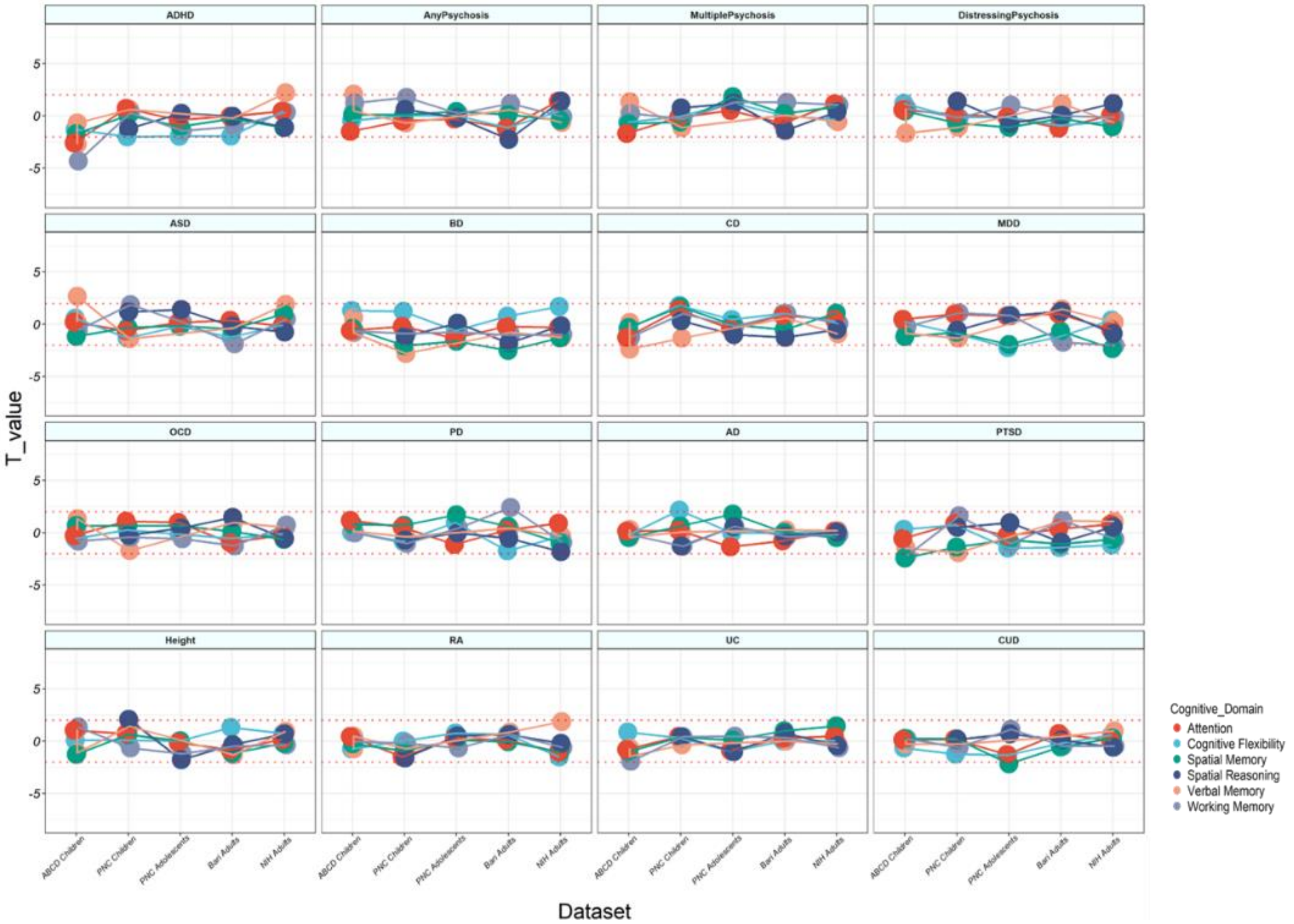


Supplementary figure 5: *cognitive performance and PGS for all other disorders in ABCD, PNC, Bari and NIH. T values for all PGS are shown on the y-axis. Disorders are attention deficit hyperactivity disorder (ADHD), Psychosis (Any /distressing experiences), autism spectrum disorder (ASD), bipolar disorder (BD), Crohn's disease (CD), Depression (MDD), obsessive compulsive disorder (OCD), Parkinson's disease (PD), Alzheimer's disease (AD), post-traumatic stress disorder (PTSD), body height (Height), rheumatoid arthritis (RA), ulcerative colitis (UC) and cannabis use disorder (CUD).*

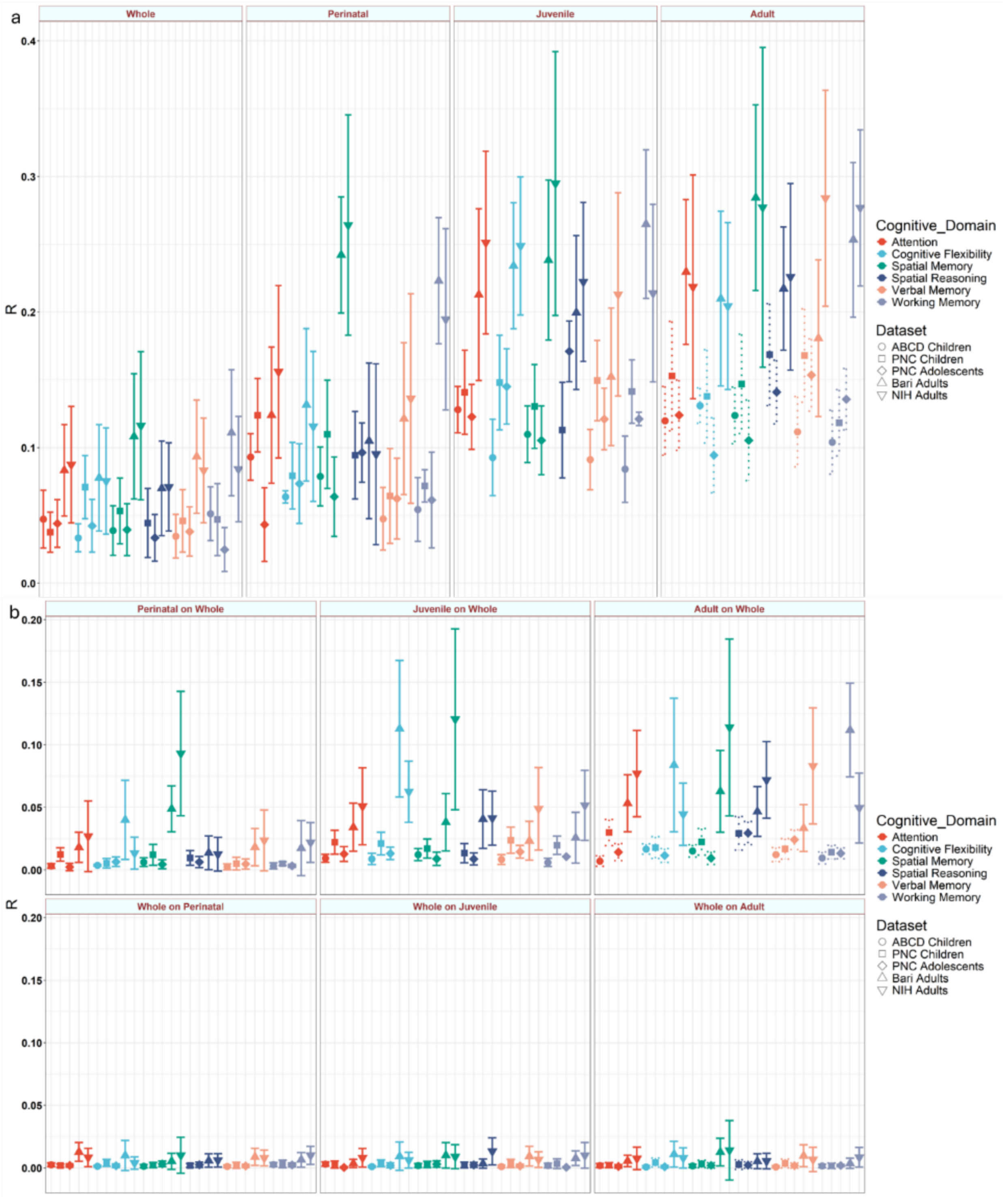


Supplementary figure 6: *a) Pearson's correlation R between observed and predicted cognitive performance with standard PGS and age-parsed PRS for SCZ risk and resilience variants. b) Variance explained ($R^2$) by parsed and full PGS models for SCZ risk and resilience variants on their statistical residuals respectively. The first row indicated the variances explained by age-parsed models (perinatal, juvenile and adult) on standard PGS residuals, while the second row indicates variance explained by the standard PGS model on age-parsed residuals.*

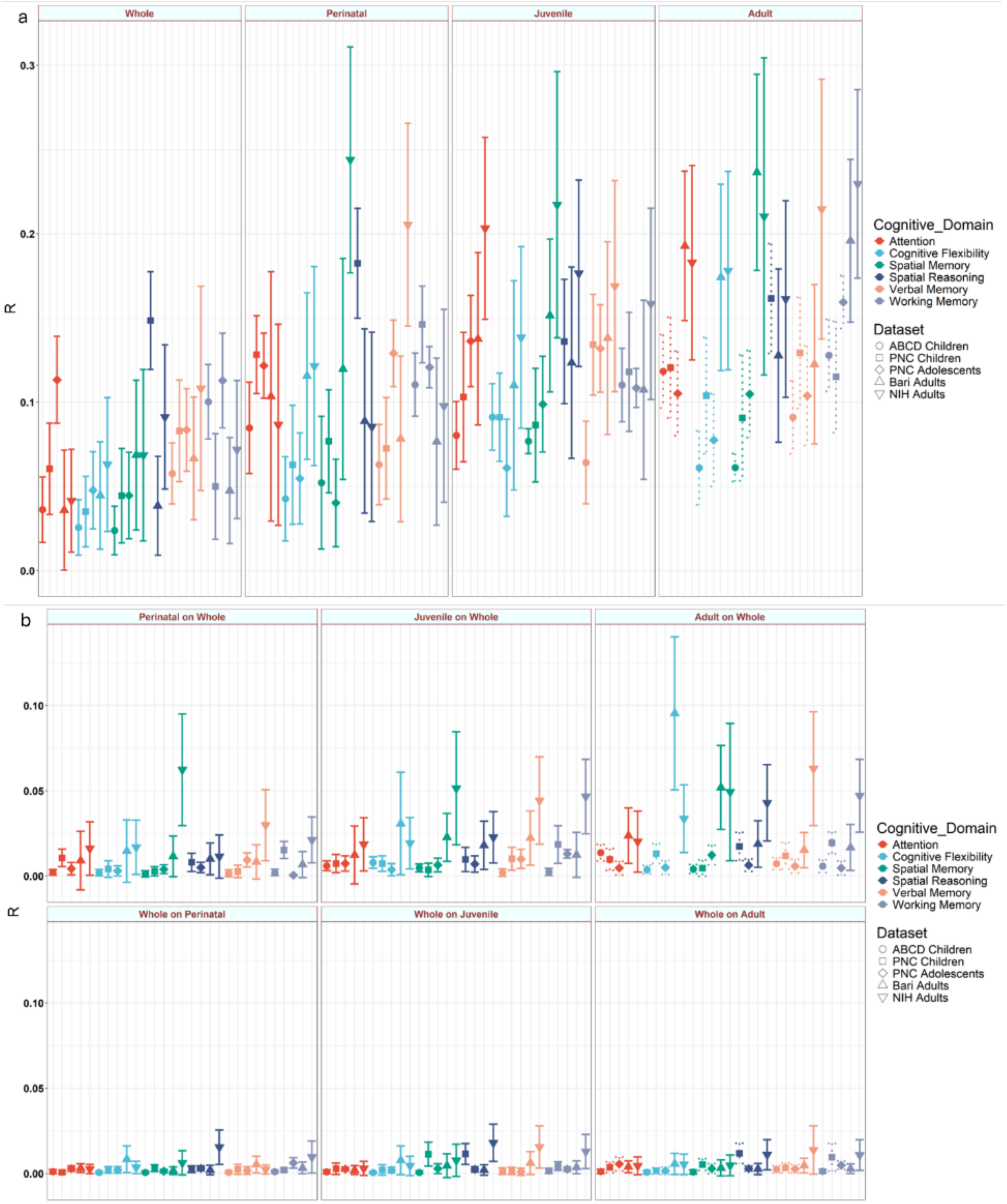


Supplementary figure 7: *a) Pearson's correlation R between observed and predicted cognitive performance with standard PGS and age-parsed PRS for cognitive intelligence variants. b) Variance explained ($R^2$) by parsed and full PGS models for cognitive intelligence variants on their statistical residuals respectively. The first row indicated the variances explained by age-parsed models (perinatal, juvenile and adult) on standard PGS residuals, while the second row indicates variance explained by the standard PGS model on age-parsed residuals.*

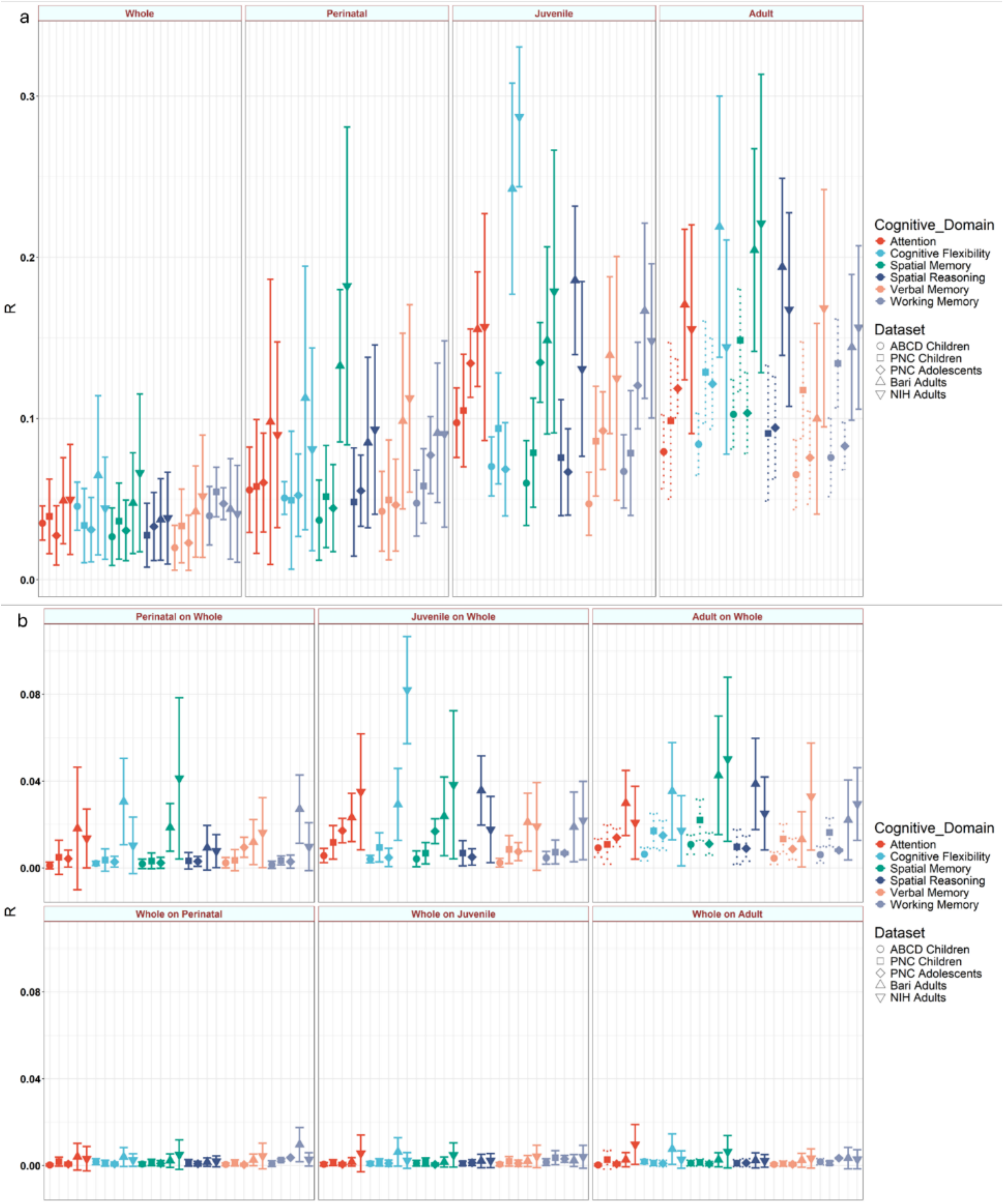


Supplementary figure 8: *a) Pearson's correlation R between observed and predicted cognitive performance with standard PGS and age-parsed PRS for general intelligence variants. b) Variance explained ($R^2$) by parsed and full PGS models for general intelligence variants on their statistical residuals respectively. The first row indicated the variances explained by age-parsed models (perinatal, juvenile and adult) on standard PGS residuals, while the second row indicates variance explained by the standard PGS model on age-parsed residuals.*

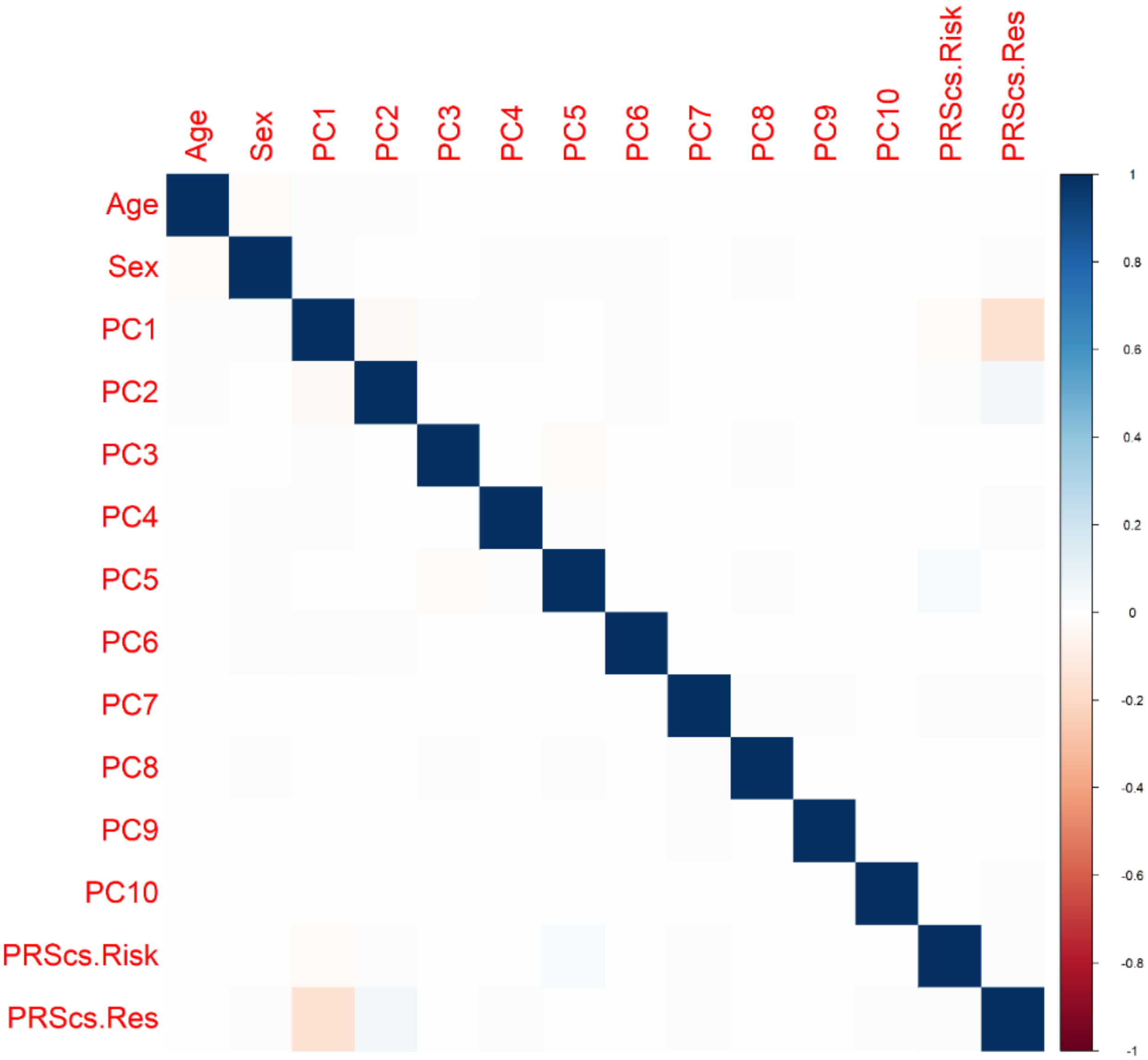


Supplementary figure 9: Correlations between each covariate used in the linear model
*Correlations between all variables, namely age, biological sex, principal components 1-10 and polygenic scores for schizophrenia risk and resilience, used within the linear model analyses. There is no significant correlation between any pair of variables.*

**Supplementary file: SynGO analysis**

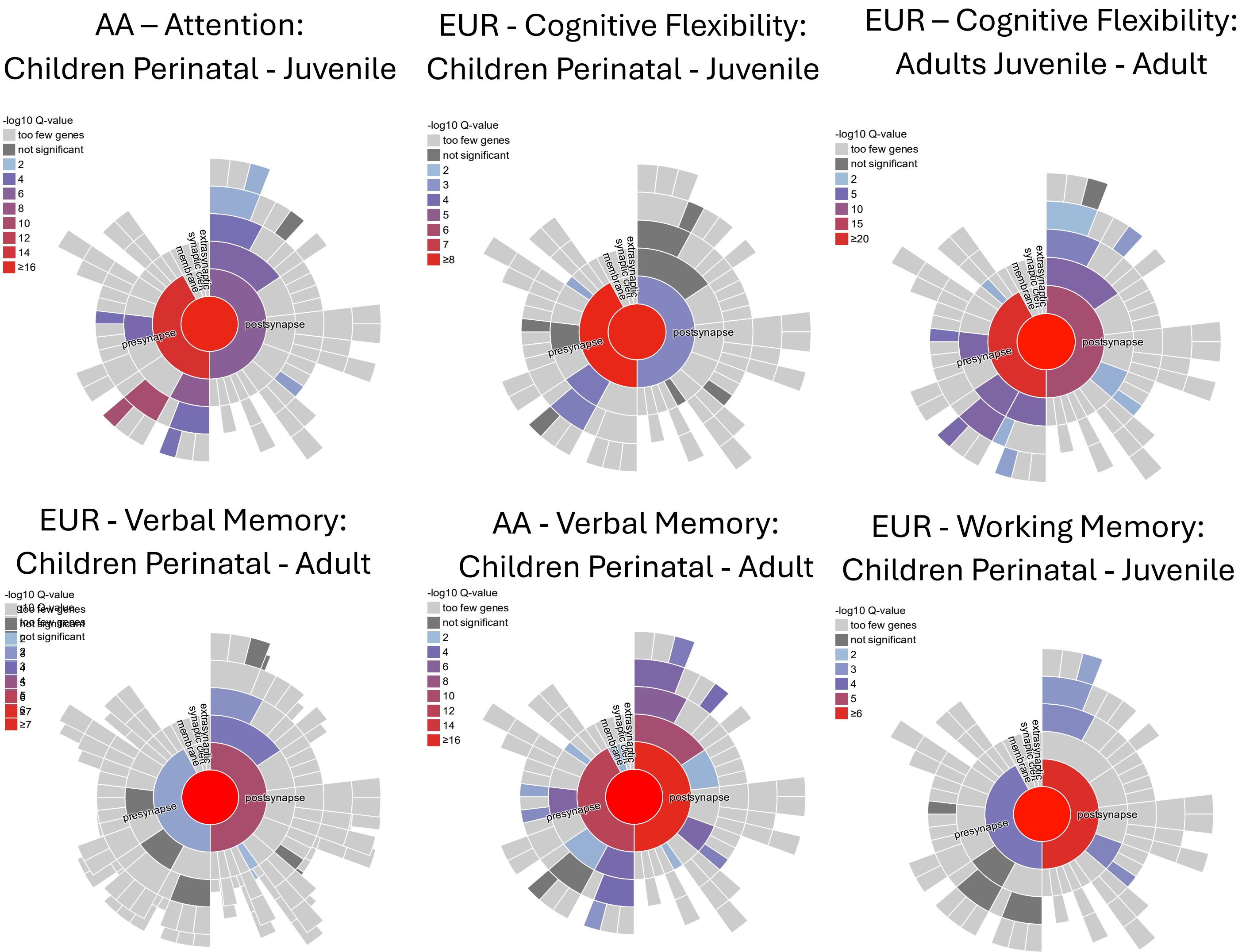


Supplementary Figure 10: SynGO analysis for overlapping gene co-expression modules within ancestry across different age stages. Attention shows overlap in AA individuals across perinatal and juvenile age stages in the children cohort. Cognitive flexibility shows an overlap of overexpressed synaptic functions across children and adult cohorts in the perinatal – juvenile and adult co-expression networks. Verbal memory shows synaptic overexpression across perinatal and adult age stages in both AA and EUR individuals. Finally, EUR samples show overlap in synaptic overexpression across perinatal and juvenile age stages in working memory tasks.

Supplementary Table 1 QC steps for genotypes

| QC STEP | THRESHOLD | COMMAND LINE | PROGRAM |
|---|---|---|---|
| **INSERTION/DELETION REMOVAL** | 100% | --snps-only --chr 1-22 | Plink |
| **GENOTYPE QUALITY** | 98% | --geno 0.02 | Plink |
| **INDIVIDUAL QUALITY** | 98% | --mind 0.02 | Plink |
| **CHECK BIOLOGICAL SEX** | 25% | --check-sex 0.25 0.75 | R / Plink |
| **REMOVE OUTLIERS FOR HETEROZYGOSITY** | 80% | --exclude range MHC/inversion region --indep-pairwise 50 5 0.2 | R / Plink |
| **REMOVE SNPS NOT IN HARDY-WEINBERG EQUILIBRIUM** | 1e-6 | --hwe 1e-6 | Plink |
| **REMOVE SNPS WITH A LOW MAF FREQUENCY** | 99% | --maf 0.01 | Plink |
| **IDENTITY BY DESCENT: EXCLUDE FIRST GRADE COUSINS AND ABOVE** | 12.5% | --missing --genome | R / Plink |
