## Supplementary Psychometric Data for "Genetic Risk and Resilience for Schizophrenia Stratified by Perinatal Gene Expression Predict Adult Cognitive Performance"

**Supplementary file: Psychometric data**

**Attention**

| Cohort | Test |
| --- | --- |
| ABCD | Flanker Task |
| PNC | Penn Continuous Performance Test (PCPT) |
| Bari | Continuous Performance Test (CPT) |
| NIH | Continuous Performance Test (CPT) |

All these assessments evaluate sustained and selective attention, focusing on the ability to maintain focus and resist distractions.

Psychometric Properties:

**Flanker Task** (ABCD)

Demonstrates sensitivity to attentional control and response inhibition.

Test-Retest Reliability: Intraclass Correlation Coefficients (ICCs) ranged from 0.438 to 0.804 for reaction times (RTs) and number of completed trials. Accuracy showed lower reliability, especially in incongruent conditions.

Validity: Test-retest reliability in children: ICC = 0.92

The Toolbox Flanker Task is an adaptation of the classic Eriksen Flanker task. It measures executive function, attention, and inhibitory control. In this task, children see five arrows in a row (or, for younger children, fish). They must identify the direction of the middle arrow (or fish), ignoring the direction of the "flanking" arrows on either side. Trials can be congruent (flankers face the same way as the middle arrow) or incongruent (flankers face the opposite way). Children respond by pressing buttons indicating the direction of the middle arrow or fish. The task provides measures of speed and accuracy, and performance typically improves with age as inhibitory control and selective attention develop. The Flanker Task showed excellent test-retest reliability in children (ICC=0.92) and expected developmental improvements.

https://doi.org/10.3758/BF03203267

https://doi.org/10.1016/j.dcn.2018.02.006

The PCPT measures vigilance and visual attention (ATT). Participants view rapidly presented horizontal and vertical line segments that form complete letters or numbers. They must press the spacebar when the stimulus forms a complete number (first half) or letter (second half). Each stimulus is shown for 300 milliseconds, with a one-second inter-stimulus interval.

Scoring: Efficiency scores derived from standardized accuracy and speed metrics.

https://doi.org/10.1037/neu0000093

**Continuous Performance Tests** (CPTs): Widely validated for assessing sustained attention and impulsivity.

Test-Retest Reliability: Good to excellent reliability with ICCs ranging from 0.62 to 0.88 across various indices.

Practice Effects: Minimal to small practice effects observed over multiple assessments.

The CPT is a widely used neuropsychological measure of sustained and selective attention. In a typical CPT, participants are asked to respond to target stimuli (e.g., letters or numbers) presented rapidly and to withhold responses to non-target stimuli. Performance is assessed based on accuracy (omission and commission errors) and reaction time variability. The CPT is sensitive to attentional deficits and has been used extensively in research on ADHD and schizophrenia. Studies (Riccio et al., 2002; Cornblatt et al., 1989) show that the CPT has good reliability and validity, and it captures both inattention and impulsivity.

https://doi.org/10.1016/S0887-6177(01)00111-1

https://doi.org/10.1016/0165-1781(88)90076-5

**Memory**

**a. Spatial Memory**

| Cohort | Test |
|---|---|
| ABCD | Picture Sequence Memory Test |
| PNC | Visual Object Learning Test |
| Bari | WMS Visual Reproduction |
| NIH | WMS Visual Reproduction |

These tests assess the ability to remember visual and spatial information, crucial for navigation and understanding spatial relationships.

Picture Sequence Memory Test : Demonstrated high test-retest reliability in adults aged 20 to 85 years.

Validity: Moderate to high correlations with other working memory and executive function measures, indicating good construct validity. Test-retest reliability in children: ICC = 0.76

The Toolbox Picture Sequence Memory Test evaluates episodic memory through a sequence of 15 pictures depicting everyday activities (e.g., working on a farm). Children view the sequence and then must reproduce it in the correct order. Unlike some memory tests, this one does not involve delayed recall or recognition phases—it purely assesses immediate memory for sequential visual information. In children and adolescents, the TPSMT demonstrated moderate reliability (ICC=0.76) and expected age-related effects, with correlations to other memory tests like the Rey Auditory Verbal Learning

Test (r~0.47). However, because the TPSMT doesn't assess delayed recall, the ABCD study also uses the RAVLT for a more complete measure of episodic memory.

https://doi.org/10.1016/j.dcn.2018.02.006

Visual object learning test

The VOLT assesses episodic memory for visual shapes (SMEM). Participants memorize 10 Euclidean shapes, followed by a recognition phase involving previously seen shapes and novel distractors. The format mirrors the PWMT but uses abstract shapes instead of words.

Scoring: Efficiency scores calculated from standardized accuracy and speed values.

https://doi.org/10.1037/neu0000093

WMS Visual Reproduction: Part of the Wechsler Memory Scale, it has established reliability and validity in measuring visual memory.

Wechsler Memory Scale (WMS) Visual Reproduction (Bari, NIH)

Reliability: The WMS is widely recognized for its strong psychometric properties, including high reliability and validity in assessing visual memory.

This subtest assesses visual memory by asking participants to reproduce geometric figures after immediate and delayed recall intervals. Scoring involves accuracy of reproduction and errors. It is sensitive to visual memory deficits seen in various clinical conditions including temporal lobe epilepsy and traumatic brain injury (Wechsler, 1997).

https://doi.org/10.1016/B978-012703570-3/50007-9

**Verbal Memory**

| Cohort | Test |
| --- | --- |
| ABCD | Rey Auditory Verbal Learning Test (RAVLT) |
| PNC | Penn Word Memory Test |
| Bari | WMS Logical Memory |
| NIH | WMS Logical Memory |

These assessments evaluate the ability to encode, store, and retrieve verbal information, essential for language comprehension and communication.

RAVLT: Widely used with reported test-retest reliability ranging from 0.55 to 0.70. Validated across various populations.

Alternate Form Reliability: Coefficients ranged from 0.60 to 0.77, indicating acceptable reliability.

Validity: Demonstrated construct validity with a bifactorial structure related to learning and episodic memory retrieval.

Although not part of the NIH Toolbox, the Rey Auditory Verbal Learning Test complements the episodic memory domain in the ABCD study. The RAVLT involves reading a list of 15 unrelated words to the child across five learning trials, followed by an interference list, immediate recall, delayed recall, and recognition. This test captures learning rate, short- and long-term recall, and recognition memory, offering a more comprehensive assessment of episodic memory compared to the TPSMT alone. It's particularly useful because it evaluates memory consolidation and retrieval, which are critical aspects of episodic memory.

https://doi.org/10.1016/j.dcn.2018.02.006

Penn Word Memory Test

The PWMT measures episodic verbal memory (VMEM). Participants are presented with 20 words (one second each) to memorize. In the recognition phase, they view 40 words (20 previously studied,

20 distractors) and indicate whether each word was seen before using a four-choice scale ("definitely not," "probably not," "probably yes," or "definitely yes").

Scoring: Accuracy and speed values are standardized (z-scores) and combined into an efficiency score (z-accuracy + z-speed).

https://doi.org/10.1037/neu0000093

WMS Logical Memory: A component of the Wechsler Memory Scale, known for its robust psychometric properties in assessing verbal memory.

This subtest evaluates verbal episodic memory by asking participants to recall short stories immediately and after a delay. It is sensitive to memory deficits in Alzheimer's disease and other neurodegenerative conditions (Wechsler, 1997).

https://doi.org/10.1016/B978-012703570-3/50007-9

**Spatial Reasoning**

| Cohort | Test |
|---|---|
| ABCD | — |
| PNC | Penn Line Orientation Test |
| Bari | Trail Making Test – Part A (TMT-A) |
| NIH | Trail Making Test – Part A (TMT-A) |

These tests measure processing speed and visuospatial abilities, reflecting the capacity to quickly and accurately process visual information.

Penn Line Orientation Test

The PLOT measures spatial reasoning (SPA). Participants see two lines on the screen and must rotate one to match the angle of the other (non-rotating) line. This task taps into spatial orientation and reasoning skills.

Scoring: Efficiency scores from standardized z-scores of accuracy and response speed.

https://doi.org/10.1037/neu0000093

TMT-A: Demonstrates good test-retest reliability and is sensitive to changes in processing speed.

Test-Retest Reliability: ICCs ranged from 0.76 to 0.89, indicating good reliability.

Validity: Sensitive to processing speed and cognitive flexibility, commonly used in assessing executive function.

The TMT-A is a measure of processing speed, visual search, scanning, and motor tracking. Participants are asked to connect numbers in order as quickly as possible. The time to complete the task is the primary measure. TMT-A has shown strong test-retest reliability and sensitivity to neurological impairment, making it a common tool in both clinical and research.

https://doi.org/10.1038/nprot.2006.390

**Executive Functions**

**Working Memory**

| Cohort | Test |
|---|---|
| ABCD | List Sorting Working Memory Test |
| PNC | Letter N-Back Test |
| Bari | N-Back Test |
| NIH | N-Back Test |

These assessments focus on the capacity to hold and manipulate information over short periods, a core component of executive functioning.

List Sorting Working Memory Test: Part of the NIH Toolbox, it has demonstrated excellent test-retest reliability and construct validity.

Test-Retest Reliability: Demonstrated excellent reliability in adults aged 20 to 85 years.

Validity: Moderate to high correlations with other working memory and executive function measures, supporting construct validity. Test-retest reliability: ICC = 0.86

The Toolbox List Sorting Working Memory Test assesses working memory by asking children to sequence items by size within categories (like animals or foods). The task starts with lists of two items and progresses up to seven items, depending on performance. In a second phase, children must sort and remember items from two different categories presented together. Scoring is based on whether the child correctly orders items from smallest to largest within the category. The TLSWMT showed good test-retest reliability (ICC=0.86) and developmental sensitivity. It also showed convergent validity with other working memory tasks (like the WISC-IV Letter-Number Sequencing) and related executive function measures.

https://doi.org/10.1016/j.dcn.2018.02.006

Penn Letter N-Back Test

The Letter N-Back Test measures working memory (WM). Participants view a continuous series of letters and must respond when the current letter matches one seen n-steps back (0-back: press for "X"; 1-back: match previous letter; 2-back: match letter two trials earlier). The inter-stimulus interval is 2.5 seconds, with each letter presented for 0.5 seconds.

Scoring: Efficiency scores created from standardized accuracy and speed z-scores.

https://doi.org/10.1037/neu0000093

N-Back Test: Shows high test-retest reliability, with Pearson coefficients ranging from 0.81 to 0.88.

Test-Retest Reliability: Pearson or Spearman coefficients ranged from 0.81 to 0.88, indicating excellent reliability.

Validity: Performance correlated significantly with cognitive assessments like the Montreal Cognitive Assessment-Basic (MoCA-BC), supporting its validity.

The N-Back is a working memory and updating task. Participants are presented with a continuous stream of stimuli (letters, numbers, images) and must indicate when the current stimulus matches the one presented "n" trials before. It measures updating, monitoring, and working memory load. Performance is scored based on accuracy and reaction time. It has been shown to activate the dorsolateral prefrontal cortex and parietal regions.

https://doi.org/10.1002/hbm.20131

**Cognitive Flexibility**

| Cohort | Test |
|---|---|
| ABCD | Dimensional Change Card Sort |
| PNC | Penn Conditional Exclusion Test |
| Bari | Wisconsin Card Sorting Test (WCST) |
| NIH | Wisconsin Card Sorting Test (WCST) |

These tests assess the ability to adapt thinking and behavior in response to changing rules or environments, a key aspect of executive functioning.

WCST: Extensively used to measure abstract reasoning and the ability to shift cognitive strategies.

Dimensional Change Card Sort

The Toolbox Dimensional Change Card Sort measures cognitive flexibility—the ability to switch between different rules or perspectives. Children sort cards by one dimension (like color), then switch to a different dimension (like shape), and finally alternate between sorting rules. It captures both accuracy and speed. The TDCCS has excellent reliability (ICC=0.92) and expected developmental patterns, with strong convergent validity with other executive function tasks. It's a classic measure of set-shifting ability—a core executive function.

Test-retest reliability in children: ICC = 0.92

https://doi.org/10.1038/nprot.2006.46

https://doi.org/10.1016/j.dcn.2018.02.006

Penn Conditional Exclusion Test

Penn Conditional Exclusion Test (PCET)

The PCET (Kurtz et al., 2004) measures executive functioning, specifically abstraction and mental flexibility. Participants are presented with four objects and must identify the one that does not belong, based on implicit rules about shape, size, or configuration. Feedback is provided ("correct"/"incorrect"), and the exclusion rule changes after ten consecutive correct responses, requiring participants to adapt to new sorting principles.

Scoring: Efficiency scores based on standardized accuracy and speed metrics.

https://doi.org/10.1037/neu0000093

Wisconsin Card Sorting Test (WCST) (Bari, NIH)

Reliability: Widely used with established reliability in assessing executive functions, particularly cognitive flexibility.

Validity: Effective in identifying frontal lobe dysfunction and assessing abstract reasoning and problem-solving abilities.

https://api.semanticscholar.org/CorpusID:65194761
